# Single cell multi-omic analysis identifies a *Tbx1*-dependent multilineage primed population in the murine cardiopharyngeal mesoderm

**DOI:** 10.1101/2020.12.24.424342

**Authors:** Hiroko Nomaru, Yang Liu, Christopher De Bono, Dario Righelli, Andrea Cirino, Wei Wang, Silvia E. Racedo, Anelisa G. Dantas, Chenleng Cai, Claudia Angelini, Lionel Christiaen, Robert G. Kelly, Antonio Baldini, Deyou Zheng, Bernice Morrow

## Abstract

The poles of the heart and branchiomeric muscles of the face and neck are formed from the cardiopharyngeal mesoderm (CPM) within the pharyngeal apparatus. The formation of the cardiac outflow tract and branchiomeric muscles are disrupted in patients with 22q11.2 deletion syndrome (22q11.2DS), due to haploinsufficiency of *TBX1*, encoding a T-box transcription factor. Here, using single cell RNA-sequencing, we identified a multilineage primed population (MLP) within the CPM, marked by the *Tbx1* lineage, which has bipotent properties to form cardiac and skeletal muscle cells. The MLPs are localized within the nascent mesoderm of the caudal lateral pharyngeal apparatus and provide a continuous source of progenitors that undergo TBX1-dependent progression towards maturation. *Tbx1* also regulates the balance between MLP maintenance and maturation while restricting ectopic non-mesodermal gene expression. We further show that TBX1 confers this balance by direct regulation of MLP enriched genes and downstream pathways, partly through altering chromatin accessibility. Our study thus uncovers a new cell population and reveals novel mechanisms by which *Tbx1* directs the development of the pharyngeal apparatus, which is profoundly altered in 22q11.2DS.

## Introduction

The heart develops from two successive waves of mesodermal progenitor cells during early embryogenesis. The first heart field (FHF) constitutes the first wave of mesodermal derived cardiac progenitors and results in the primitive beating heart, while the second heart field (SHF) forms the second wave that builds upon the two poles of the heart^1,2^. The SHF can be anatomically partitioned to the anterior SHF (aSHF; ^3,4^) and posterior SHF (pSHF; ^4–6^), whose cells migrate to the heart via the outflow tract or inflow tract, respectively. Expression of *Mesp1* at gastrulation marks the earliest mesodermal cells that will form the heart^7^. Using single cell RNA-sequencing (scRNA-seq) of the *Mesp1* lineage, it was discovered that the FHF, aSHF and pSHF are specified at gastrulation^8^.

Retrospective clonal analysis^9,10^ and lineage tracing studies^11^ revealed that the branchiomeric skeletal muscles (BrM) of the craniofacial region and neck share a clonal relationship with the SHF. The bipotent nature of these cardiac and skeletal muscle progenitor cells is supported by studies of the ascidian, *Ciona,* an invertebrate chordate, in which single cells gives rise to both cardiac and skeletal muscle cells^12^. When taken together, a new term, the cardiopharyngeal mesoderm (CPM), was introduced to clearly include both SHF cardiac and skeletal muscle progenitor populations^2^. A cartoon of these populations is shown in Fig. 1a. The *Tbx1* gene, encoding a T-box transcription factor, and gene haploinsufficient in 22q11.2 deletion syndrome (22q11.2DS), is expressed in the CPM and is required for cardiac outflow tract and BrM development^2^, implicating its essential roles in the CPM.

**Figure 1.**
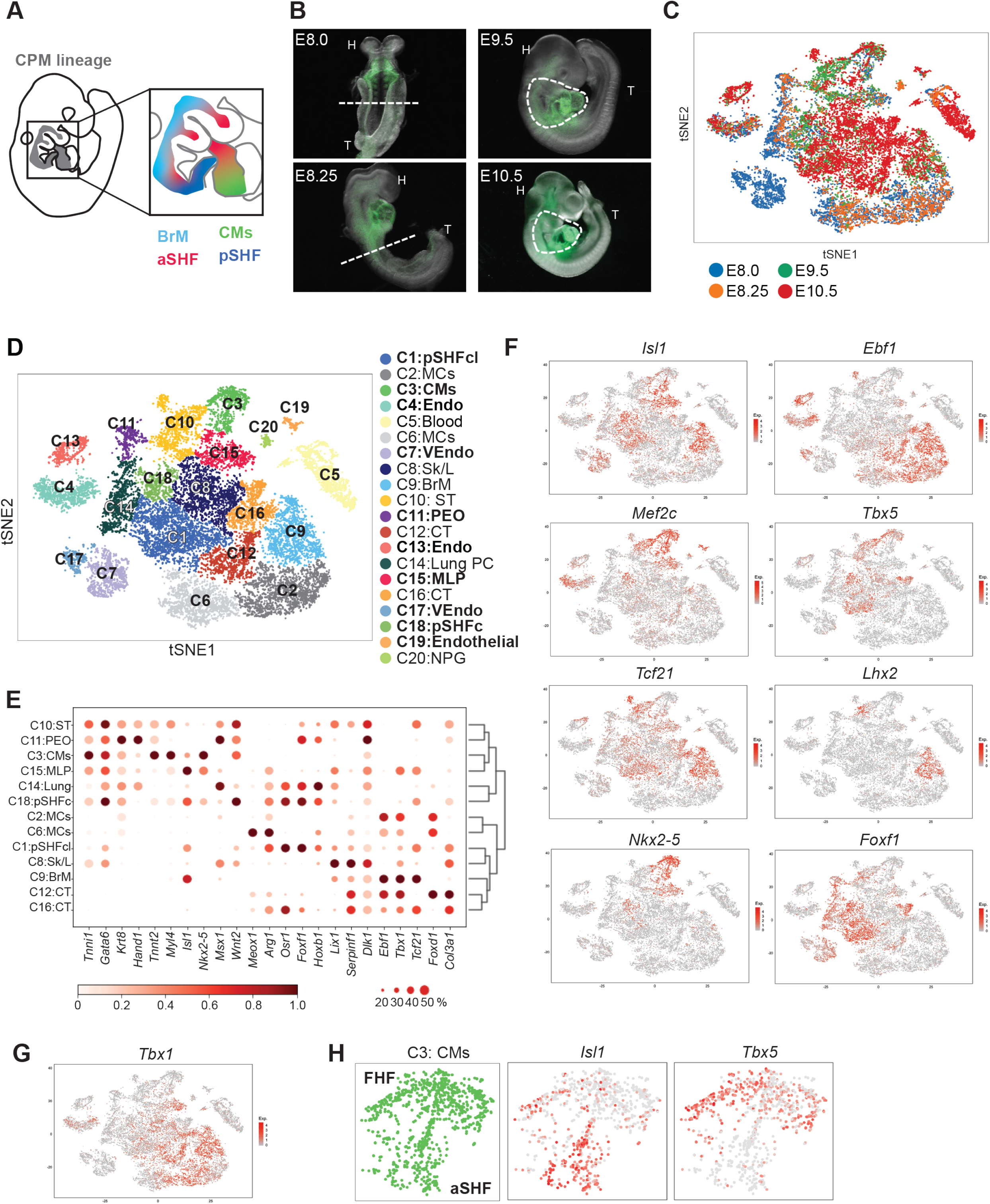
Single-cell analysis of *Mesp1*+ lineages at E8.0 to E10.5 identifies CPM lineages. **A.** Cartoon of an E9.5 embryo in a right lateral view. The CPM is colored in gray. Components of the CPM include the branchiomeric muscle progenitor cells (BrM; aqua), anterior SHF (aSHF; red) and part of the posterior SHF (pSHF; blue). Because cardiomyocyte progenitors (CMs) that are shown in green are from both the FHF and CPM, only CPM-derived CMs are included. **B.** Whole embryo images with GFP fluorescence of *Mesp1*^*Cre*^; *ROSA26-GFP*^*flox*/+^ embryos at E8.0, E8.25, E9.5 and E10.5 for scRNA-seq. White dotted line represents region that was dissected. The upper half of the body was collected at E8.0 and E8.25. The pharyngeal apparatus and heart were collected at E9.5 and E10.5. Only fluorescence sorted GFP positive cells were used for scRNA-seq. Abbreviations: head (H), Tail (T). **C.** t-distributed stochastic neighbor embedding (tSNE) plot colored by the developmental stage of the samples. **D.** tSNE plot colored by clusters. 20 clusters were identified based on gene expression patterns. Cardiac relevant clusters are in bold font. C1: pSHFcl, posterior CPM includes cardiac and lung progenitor cells; C2: MCs, mesenchyme expressing epithelial-mesenchymal transition markers; C3: CMs, cardiomyocyte progenitor cells; C4: Endo, endocardium and endothelial cells; C5: Blood, blood cells; C6: MCs, mesenchyme expressing epithelial-mesenchymal transition markers; C7: VEndo, vascular endothelial cells; C8: Sk/L, skeleton/limb progenitor cells; C9: BrM, branchiomeric muscle progenitor cells; C10: ST, septum transversum progenitor cells; C11: PEO, proepicardial organ; C12: CT, connective tissue progenitor cells; C13: Endo, endothelial cells; C14: Lung PC, lung progenitor cells; C15: MLP, multilineage progenitor cells; C16: CT, connective tissue progenitor cells; C17: VEndo, vascular endothelial cells; C18: pSHFc, posterior CPM of cardiac progenitor cells; C19: Endothelial cells; C20: NPG, neural progenitor cells. **E.** Heatmap of average gene expression of marker genes in representative clusters. Dot size indicates the fraction of cells expressing the genes in each cluster and color indicates scaled mean expression. **F.** tSNE plots for showing the expression of the CPM marker genes. The color spectrum from grey to red indicates expression level from low to high. **G.** tSNE plot for showing the expression level of *Tbx1*. The color spectrum from grey to red indicates expression levels from low to high. **H.** tSNE plots of CMs showing the expression level of CM and CPM marker genes. *Tbx5*+; *Isl1*- cells identify the FHF population, and *Tbx5*-; *Isl1*+ cells for the aSHF population.

A total of 60-75% of patients with 22q11.2DS have cardiac outflow tract defects, which often require life-saving surgery during the neonatal period ^13^. Additionally, most individuals with this condition have speech, feeding and swallowing difficulties in infancy, due in part to BrM hypotonia^14^. Further, heterozygous mutations in rare, non-deleted individuals, phenocopy the symptoms of the deletion^15^. Inactivation of *Tbx1* in the mouse results in a persistent truncus arteriosus (PTA)^16–18^ and significant failure to form the BrMs^19^. Although there are many studies of *Tbx1*, we do not yet understand its functions on a single cell level, which is needed to elucidate the true molecular pathogenesis of 22q11.2DS.

The CPM is distributed throughout the embryonic pharyngeal apparatus during early gestation. The pharyngeal apparatus consists of individual bulges of cells termed arches that form in a rostral to caudal manner from mouse embryonic days (E)8-10.5. The cellular and molecular mechanisms of how CPM cells in the pharyngeal apparatus both are maintained in a progenitor state and are allocated to form the heart and BrMs in mammals, are unknown.

To fill these gaps, we performed scRNA-seq of the CPM at multiple stages during embryogenesis. We discovered a multilineage primed progenitor (MLP) population within the CPM, which is maintained from E8-10.5 and has differentiation branches toward cardiac and skeletal muscle fates, serving as common lineage progenitors. The MLP cells are localized to the nascent lateral mesoderm of the pharyngeal apparatus, deploying cells to the heart and BrMs. We found that the *Tbx1* cell lineage marks the MLPs and TBX1 activity is critical for their function. Inactivation of *Tbx1* disrupts MLP lineage progression and results in ectopic expression of non-mesoderm genes. We further identify the gene regulatory network downstream of *Tbx1* in the MLPs providing insights into the molecular mechanism of mammalian CPM function, essential for understanding the etiology of 22q11.2DS.

## Results

### Identification of common progenitor cells in the CPM

To identify the various populations that constitute the CPM (Fig. 1a), we performed droplet-based scRNA-seq on fluorescence activated cell sorted GFP expressing cells from microdissected *Mesp1^Cre^; ROSA26-GFP*^*flox*/+^ embryos at E8.0, E8.25, E9.5 and E10.5 (Table 1, Fig. 1b; Extended Data Fig. 1a, b). These stages were chosen because they are the critical periods when *Tbx1* is expressed and when the pharyngeal apparatus is dynamically elongating; this is coordinated with heart development and BrM specification. To better understand the developmental connection, we integrated the four time point datasets with Scran software^20^ (Fig. 1c) and identified 20 cell clusters using densityClust v0.3^21^ (Fig. 1d, e, Extended Data Fig. 1c, e, and Supplementary Table 1). Utilizing knowledge of specific gene expression in each cluster (Fig. 1e, Extended Data Fig. 1c, e), we identified cell types of all the clusters, half of which include cardiovascular progenitor cell populations or their derivatives (Fig. 1d; bold font).

**Table1.** Summary of scRNA-seq samples.

| Time point | No. of embryos | somites | cell viability | cell number (submitted) | cell number (captured) | Mean reads/cell | Median genes /cell | Tissue |
| --- | --- | --- | --- | --- | --- | --- | --- | --- |
| *Mesp1<sup>Cre</sup>; ROSA26-GFP<sup>flox/+</sup>* |  |  |  |  |  |  |  |  |
| E8.0 | 7 | 3-4 | 90% | 16,416 | 7,581 | 58,422 | 4,380 | Half of the body |
| E8.25 | 5 | 6-7 | 95% | 8,364 | 2,956 | 151,357 | 5,363 | Half of the body |
| E9.5 | 6 | 19-22 | 80% | 12,000 | 4,055 | 80,919 | 4,036 | PA and heart |
| E10.5 | 4 | 30-31 | 80% | 18,038 | 8,070 | 52,326 | 3,673 | PA and heart |
| *Mesp1<sup>Cre</sup>; ROSA26-GFP<sup>flox/+</sup>; Tbx1<sup>flox/flox</sup>* |  |  |  |  |  |  |  |  |
| E9.5 | 1 | 22 | 83% | 13,788 | 5,157 | 65,997 | 5,015 | PA and heart |
| *Tbx1<sup>Cre/+</sup>; ROSA26-GFP<sup>flox/+</sup>* |  |  |  |  |  |  |  |  |
| E8.5 | 7 | 8-9 | 88% | 4,975 | 1,434 | 136,347 | 3,683 | Half of the body |
| E9.5 | 10 | 19-21 | 87% | 18,300 | 5,453 | 54,317 | 3,994 | PA and heart with back |
| *Tbx1<sup>Cre/flox</sup>; ROSA26-GFP<sup>flox/+</sup>* |  |  |  |  |  |  |  |  |
| E8.5 | 4 | 9-10 | 92% | 2,820 | 1,090 | 140,352 | 4,374 | Half of the body |
| E9.5 | 3 | 19-22 | 97% | 11,560 | 6,267 | 54,892 | 3,561 | PA and heart with back |

Both aSHF and pSHF contribute to the cardiac outflow tract, while only pSHF cells form the inflow tract^22,23^. Marker genes for the CPM include *Tbx1*, *Isl1* and *Tcf21*, among other genes^1,24^ (Fig. 1f, g). Cluster C9 contains BrM progenitor cells identified by expression of *Tcf21, Lhx2* and *Myf5* ^24^ (Fig. 1f and Extended Data Fig. 1e). Clusters C1 and C18 contains pSHF populations as identified by expression of *Hoxb1* ^5^, *Tbx5, Foxf1,* and *Wnt2*^25,26^ (Fig. 1f and Extended Data Fig. 1e). Many of the pSHF cells are located more medially and caudally in the embryo and also contribute to posterior organ development, such as the formation of the lung^25,26^. The cells in C3 express *Nkx2-5* and *Mef2c*^27^ and cardiac structural protein genes (*Tnnt2, Tnni1, Myl4*, (Fig. 1f; Extended Data Fig. 1e). In addition, subdomains express either FHF genes (*Tbx5* but not *Isl1* or *Tcf21*) or aSHF genes (*Fgf10*, *Isl1, Tcf21* but not *Tbx5* (Fig. 1h). We discovered that cluster C15 expresses genes shared by the CPM clusters, including *Tbx1, Isl1, Mef2c, Tcf21* and *Foxf1* (Fig. 1f). Based upon this, we refer C15 as the common multilineage progenitors within the CPM. This shows for the first time that CPM progenitors can be distinguished from more mature CPM states by their multilineage primed gene signatures.

### Multilineage progenitors of the CPM differentiate to cardiac and skeletal muscles

We next investigated the relationship between the common lineage progenitors and more differentiated CPM cells using partition-based graph abstraction (PAGA)^28^. Several clusters that are not part of the CPM and were already well separated in the above cluster analysis (C4, C5, C7, C13 and C17), were excluded from PAGA analysis (Extended Data Fig. 1d). The PAGA analysis partitioned the CPM cells into six branches (Fig. 2a), connecting all the mesodermal cell populations (Fig. 1c). Convergent results from pseudotime analysis (Fig. 2b, Extended Data Fig 2b) and real time point information (Fig. 2c, Extended Data Fig 2a), infer that cells in C15 in the center are in a progenitor state of the CPM while more differentiated cells are towards the outside in each branch (BrM, C9; cardiomyocytes [CMs] with aSHF, C3; part of the pSHF, C1+C18; Fig. 1d).

**Figure 2.**
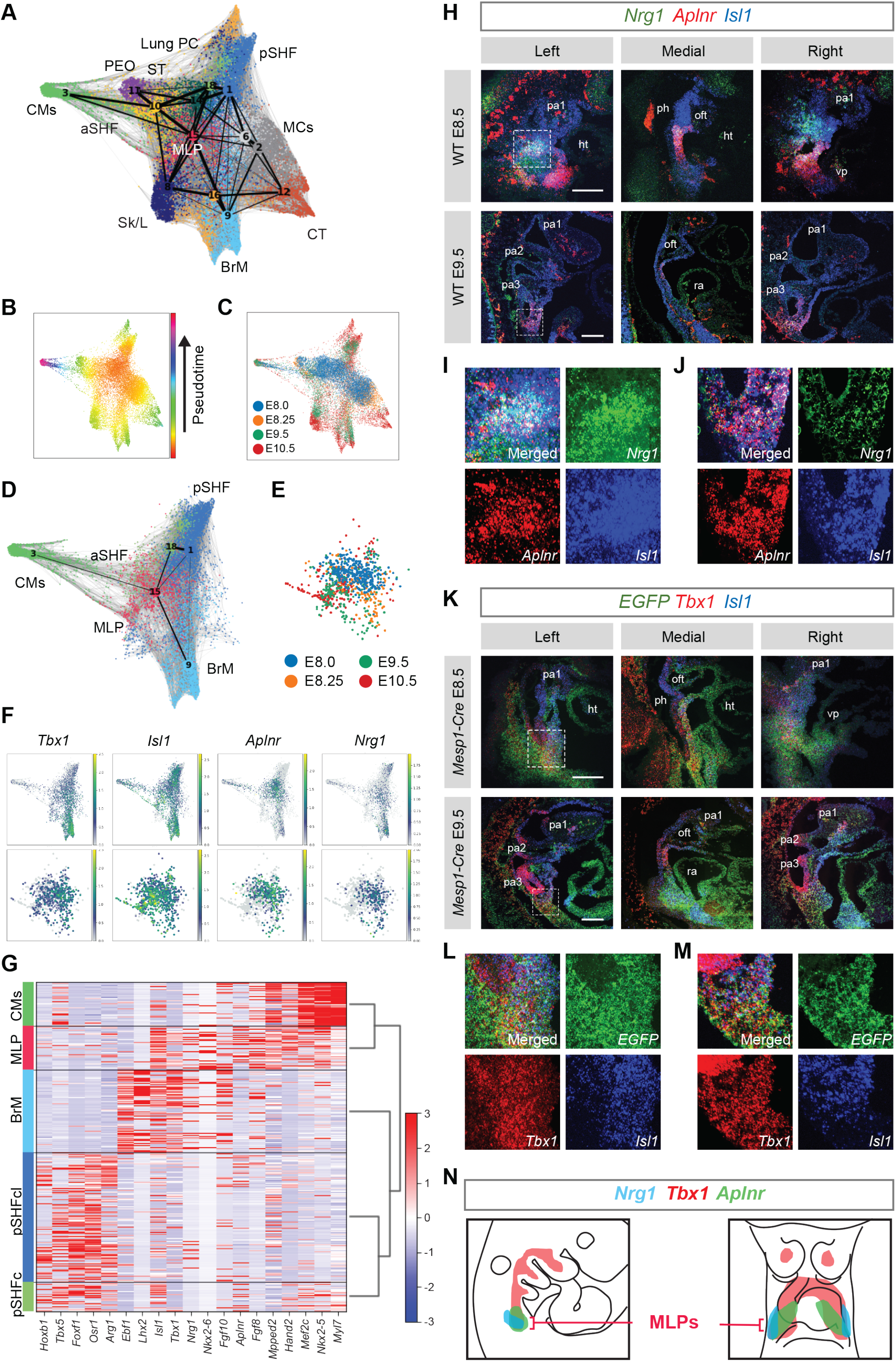
MLPs are localized to the caudal pharyngeal apparatus. **A.** Single-cell embedding graph colored by clusters along with a PAGA (Partition-based graph abstraction generates) plot. Cluster numbers are consistent with Figure 1C. CMs (C3); PEO (C11); ST (C10); Lung PC (C14); pSHF (C1, C18), pSHF; MCs (C2, C6); CT (C16); BrM (C9); Sk/L (C8); aSHF; MLP (C15), in the center of the graph; see Figure 1 for definitions. **B.** Single-cell embedding graph colored by pseudotime. The color spectrum from red/orange is early pseudotime points to blue/purple as late pseudotime points. **C.** Single-cell embedding graph colored by the developmental stage of the samples. **D.** Single-cell embedding graph with PAGA plot of the CPM. pSHF (C1, C18), BrM (C9), CMs (C3) and MLP (C15) clusters are included. **E.** Single-cell embedding graph of only MLPs colored by the stage of the samples. **F.** Single-cell embedding graph of the CPM (above) and MLPs (below) colored by gene expression level. The color spectrum from blue, then green to yellow indicates expression levels from low to high. Grey indicates no expression. **G.** Heatmap of expression of genes enriched in the CPM. Row indicates the expression of each cell. **H.** RNAscope *in situ* hybridization with *Nrg1*, *Aplnr* and *Isl1* mRNA probes on sagittal sections from wild-type embryos at E8.5 (upper panel; n = 3) and E9.5 (lower panel; n = 3). Green, *Nrg1*; Red, *Aplnr*; Blue, *Isl1*. White dotted line indicates the position in images with higher magnification (F and G). Scale bar; 100 μm. **I** and **J.** Higher magnification image of RNAscope *in situ* hybridization with *Nrg1*, *Aplnr* and *Isl1* mRNA probes on sagittal section from wild-type embryo at E8.5 (F) and E9.5 (G). Upper left, merged image; Upper right, green channel (*Nrg1*); Lower left, red channel (*Aplnr*); Lower right, blue channel (*Isl1*). **K.** RNAscope *in situ* hybridization with *EGFP* mRNA probe, marking *Mesp1*+ lineage cells, *Tbx1* and *Isl1* mRNA probes on sagittal sections from *Mesp1*^*Cre*^; *ROSA26-GFP*^*flox*/+^ embryos at E8.5 (upper panel; n = 3) and E9.5 (lower panel; n = 3). Green, *EGFP*; Red, *Tbx1*, Blue, *Isl1*. White dotted line indicates the position in images with higher magnification (I and J). Scale bar; 100 μm. **L** and **M.** Higher magnification image of RNAscope *in situ* hybridization with *EGFP*, *Tbx1* and *Isl1* mRNA probes on sagittal sections from *Mesp1^Cre^; ROSA26-GFP*^*flox*/+^ embryos at E8.5 and E9.5. Upper left, merged image; Upper right, green channel (*EGFP*); Lower left, red channel (*Tbx1*), Lower right, blue channel (*Isl1*). **N.** Summary of expression pattern of selected marker genes. Left, lateral view; Right, ventral view. *Nrg1* (aqua) and *Aplnr* (green) are expressed in the lateral caudal pharyngeal mesoderm cells, in which *Tbx1* (dark pink) and *Isl1* are also expressed. The region of overlap is the MLPs. Abbreviations: heart (ht), outflow tract (oft), pharyngeal arch (pa), pharynx (ph), right ventricle (RA), venous pole (vp), 1, 2 and 3 indicate the first, second and third pharyngeal arches, respectively.

Next, the gene expression in CPM progenitor cells (C15) were examined from E8-E10.5 (Fig. 2e, Extended Data Fig 2c). We searched for marker genes that are enriched in expression in MLPs, and we identified two newly appreciated genes, *Aplnr* (Apelin receptor) and *Nrg1* (Neuregulin 1) (Fig. 2f, 2g, Extended Data Fig 2d). *Aplnr* is expressed in the CPM^29^ but not known for MLPs, while *Nrg1* is not known to be a CPM gene, and it is required in the embryonic heart for the development of the chamber myocardium^30^. We examined the co-expression of genes in the cells in CPM populations. The heatmap in Fig. 2g (Extended Data Fig. 2e, f) shows expression of genes enriched in C15, with the same genes also expressed in more differentiated CPM populations, indicating that they are multilineage primed. We found that even at E10.5, these cells are still present and retain a multilineage state. Taken together, we refer to the cells belonging to C15 as the multilineage progenitors (MLPs).

### The MLPs are bilaterally localized to the caudal pharyngeal apparatus

To elucidate whether MLPs are dispersed or localized within a defined embryonic region in the pharyngeal apparatus, we performed RNAscope *in situ* hybridization analysis using probes enriched in MLPs including *Tbx1*, *Isl1*, *Aplnr* and *Nrg1* with the *Mesp1*+ lineage marked by GFP expression (Fig. 2h-m). The pharyngeal arches form in a rostral to caudal manner in which the most caudal and lateral mesoderm is the least differentiated, while the rostral mesoderm has already migrated to the core of the arch to form BrM progenitor cells or towards the poles of the heart^31,32^. In both E8.5 and E9.5 embryos, *Nrg1* and *Aplnr* co-expressing cells were found bilaterally in the lateral part of the caudal pharyngeal apparatus containing nascent mesoderm that is not yet differentiated to cardiac or skeletal muscle (Fig. 2f, h, Extended Data Fig. 2g, i). At E8.5, *Nrg1* and *Aplnr* were expressed in these regions within the forming second arch (Fig. 2i) and at E9.5, by the forming fourth arch (Fig. 2j), both overlapping with *Isl1* expression. The *Mesp1*+ lineage is marked with *GFP* expression in the second arch at E8.5 and the fourth arch at E9.5 (Fig. 2k, Extended Data Fig 2h, j). *Tbx1* and *Isl1* were also expressed in those regions (Fig. 2l, m). We suggest that the MLPs remain in the same region of the caudal pharyngeal apparatus, while they deploy cells rostrally, medially and dorsally thereby explaining in part the mechanism for the extension of the pharyngeal apparatus caudally (Fig. 2n).

### MLPs dynamically transition over time

An important question is whether MLPs as CPM progenitors, maintain the same state based upon gene expression over time. To address this, we examined differentially expressed genes in MLPs from E8-10.5. We identified core CPM genes that are expressed similarly at all time points, including *Isl1*, *Mef2c* and *Nkx2-5* (Fig. 3a, d). However, we also found that early expressing genes such as *Aplnr, Nrg1, Irx1-5*, *Fgf8/10* and *Tbx1* (Fig. 3b, e) are reduced over time, with increasing expression of cardiac developmental genes such as *Hand2, Gata3/5/6, Bmp4* and *Sema3c* (Fig 3c, f). This is consistent with the model that the MLPs continuously allocate progenitor cells to BrMs and CMs, while showing some maturation themselves (Fig. 3g). We next tested if *Tbx1* has a specific role in MLPs.

**Figure 3.**
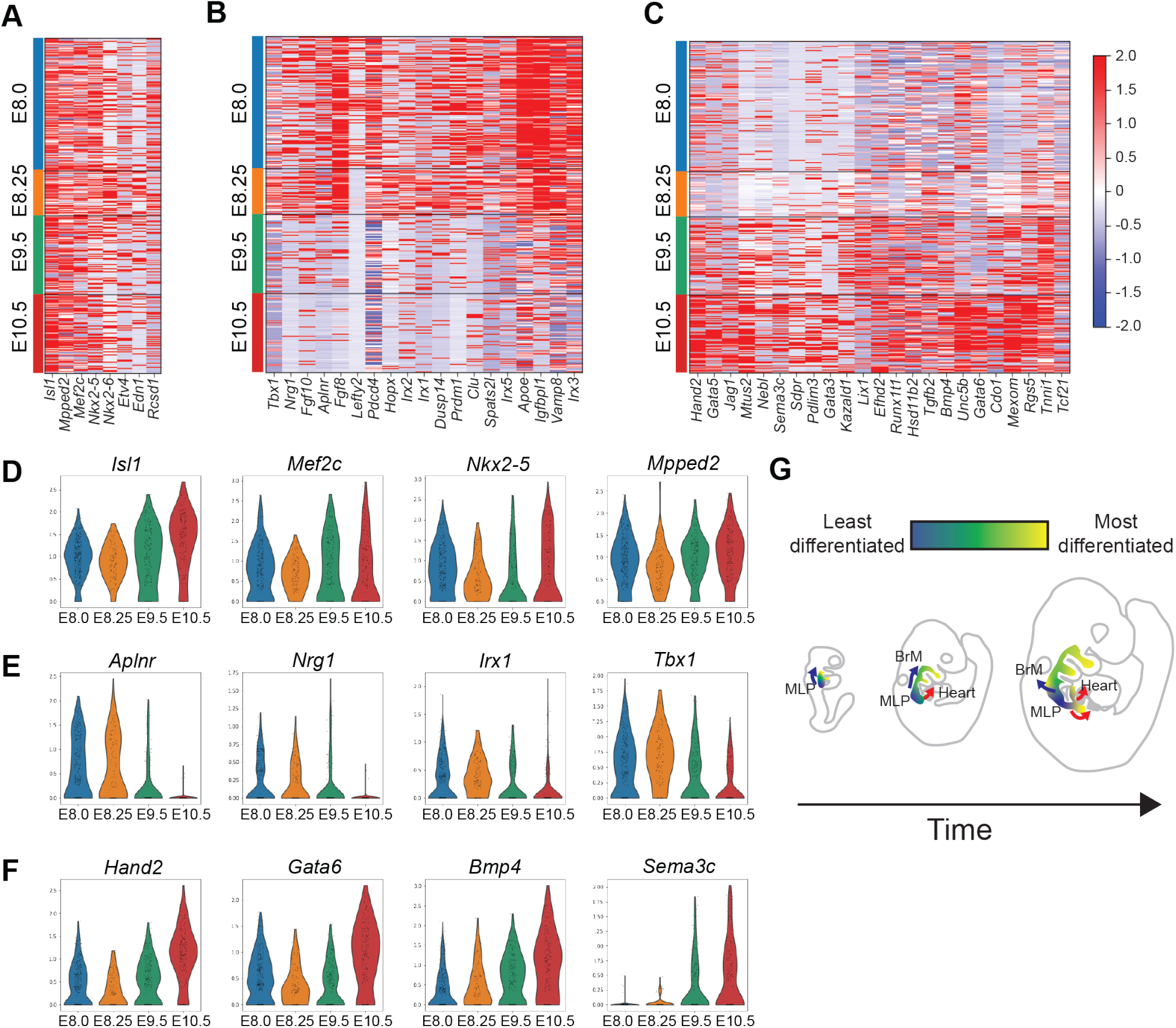
MLPs transition as cells are allocated to more differentiated states over time. **A.** Heatmap of expression of core genes in MLPs. Row indicates the expression of each cell. **B.** Heatmap of expression of the genes in earlier stage-MLPs (E8, E8.25). Row indicates the expression of each cell. **C.** Heatmap of expression of the genes in late stage-MLPs (E9.5, E10.5). Row indicates the expression in each cell. **D.** Violin plots of the expression of core genes (*Isl1, Mef2c, Nkx2-5, Mpped2*) in MLPs over time. **E.** Violin plots of expression of early-MLP genes (*Aplnr, Nrg1, Irx1, Tbx1*) in MLPs over time. **F.** Violin plots of expression of late-MLP genes (*Hand2, Gata6, Bmp4, Sema3c*) in MLPs over time. **G.** Cartoon of MLP transitions and cell allocation over time. The MLPs in the nascent pharyngeal mesoderm migrate dorsally and will differentiate to BrMs or ventrally and medially to CMs. Blue arrow indicates migration to form BrMs and red arrow(s) indicates migration to the poles of the heart to form CMs. The color spectrum from blue to yellow indicates differentiation from MLPs to their derivative cell types.

### Loss of *Tbx1* increases the proportion of MLPs to those of more differentiated states

*Tbx1* is not one of the highest expressed genes in MLPs (Figs. 1e, 2g and 3b). Yet, based upon its function in embryogenesis, we suspect it has a critical role in MLPs because its inactivation results in failed heart and BrM development. We therefore examined the *Mesp1* versus *Tbx1* lineages in control embryos to understand how the CPM lineages compare. We tested whether the *Tbx1* lineage includes the MLPs and how inactivation would affect MLP function. To address these questions, we compared embryos that were *Mesp1^Cre^;Tbx1*^+/+^ (*Mesp1^Cre^;Tbx1* Ctrl) vs *Mesp1^Cre^;Tbx1^flox/flox^* (*Mesp1^Cre^;Tbx1* cKO) at E9.5 (Fig. 4a, Table 1) and *Tbx1*^*Cre*/+^ (*Tbx1^Cre^;Tbx1* Ctrl) vs *Tbx1^Cre/flox^* (*Tbx1^Cre^;Tbx1* cKO) at E8.5 and E9.5 (Fig. 4b, Table 1). In the SwissWebster background, *Tbx1*^*Cre*/+^ and *Tbx1*^+/−^ heterozygous mice have no heart or aortic arch defects^33^ and thus serve as controls.

**Figure 4.**
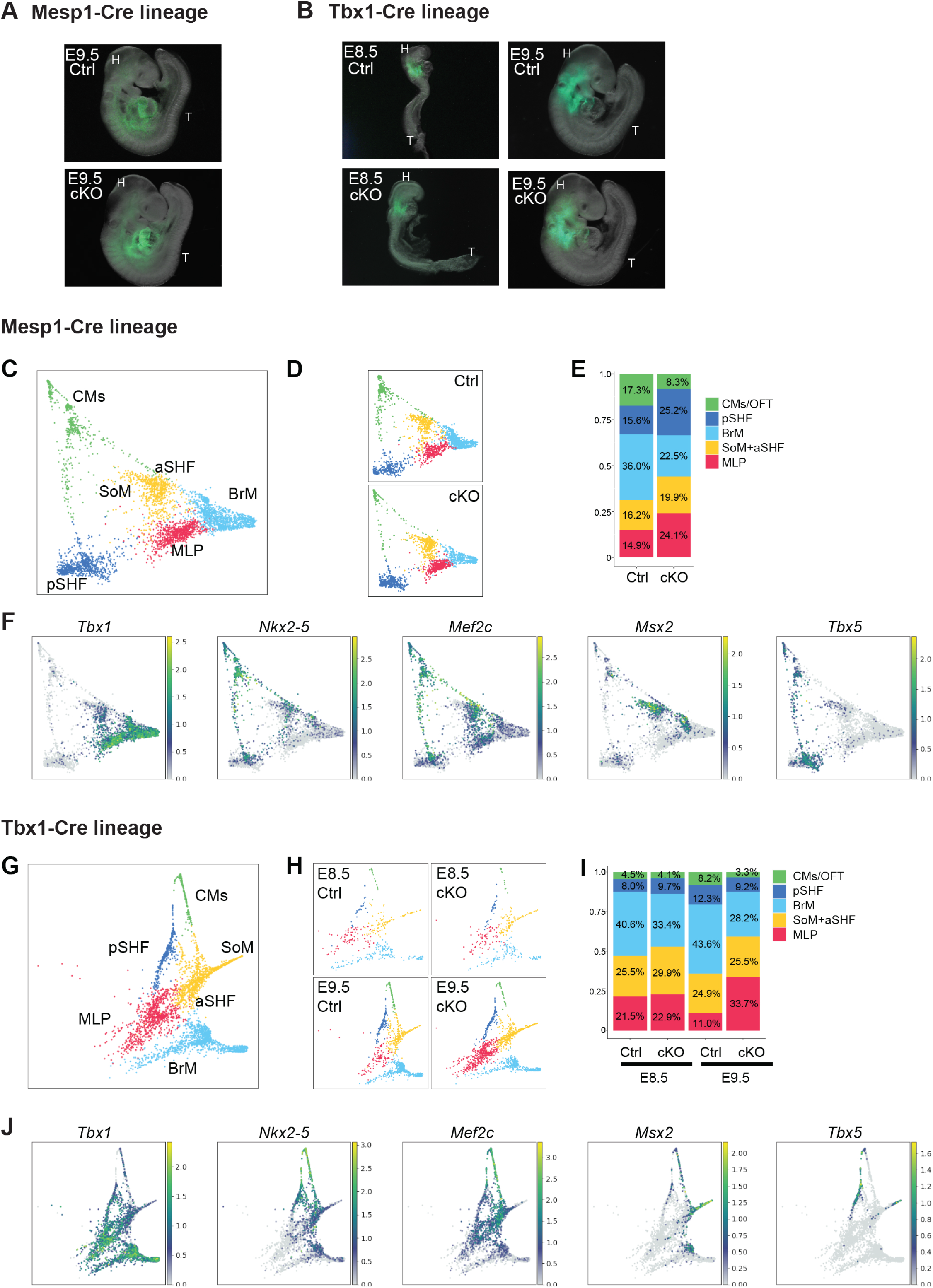
*Tbx1* is required for progression of MLPs to more differentiated states. **A.** Whole mount embryo images with GFP fluorescence of *Mesp1^Cre^; ROSA26-GFP*^*flox*/+^ (Ctrl; upper panel, similar to Fig. 1A) and *Mesp1^Cre^; ROSA26-GFP*^*flox*/+^;*Tbx1^flox/flox^* (cKO; lower panel) embryos at E9.5 used for scRNA-seq. The pharyngeal apparatus and heart were collected. The sorted GFP positive cells were used for scRNA-seq. Abbreviations: head (H), Tail (T). **B.** Whole embryo images with GFP fluorescence of *Tbx1*^*Cre*/+^; *ROSA26-GFP*^*flox*/+^ (Ctrl; upper panels) and *Tbx1^Cre/flox^; ROSA26-GFP*^*flox*/+^ (cKO; lower panels) embryos at E8.5 (left) and E9.5 (right) used for scRNA-seq. The upper half of the body was collected at E8.5. The pharyngeal apparatus and heart were collected at E9.5. Sorted GFP positive cells were used for scRNA-seq. **C.** Single-cell embedding graph of the CPM populations from *Mesp1^Cre^;Tbx1* Ctrl vs cKO datasets colored by clusters. Definitions of clusters are the same as in Figure 1, with the addition of SoM, somatic mesoderm. **D.** Single-cell embedding graph separated by genotype. Upper panel, *Mesp1^Cre^;Tbx1* Ctrl dataset. Lower panel; *Mesp1^Cre^;Tbx1* cKO dataset. **E.** The ratio of cell populations in CPM lineages in *Mesp1^Cre^;Tbx1* Ctrl and cKO datasets. A two proportion Z-test was performed in each cluster (MLP: P = 2.63e^−12^, aSHF+SoM: P=3.43e^−3^, BrM: P < 2.2e^−16^, pSHF: P = 1.12e^−12^, CM: P = 6.81e^−16^). **F.** Single-cell embedding graph of the CPM lineages from *Mesp1^Cre^;Tbx1* Ctrl and cKO datasets colored by expression level. The color spectrum from blue to yellow indicates expression level from low to high. Grey indicates no expression. **G.** Single-cell embedding graph of CPM lineages from *Tbx1^Cre^;Tbx1* Ctrl and cKO datasets at E8.5 and E9.5 datasets colored by cell clusters. **H.** Single-cell embedding graph separated by samples. Upper left, *Tbx1^Cre^;Tbx1* Ctrl dataset; Upper right, *Tbx1^Cre^;Tbx1* cKO dataset at E8.5. Lower left, *Tbx1^Cre^;Tbx1* Ctrl dataset; Lower right, *Tbx1^Cre^;Tbx1* cKO dataset at E9.5. **I.** The ratio of cell populations in CPM lineages in *Tbx1^Cre^;Tbx1* Ctrl and *Tbx1^Cre^;Tbx1* cKO datasets at E8.5 and E9.5. A two proportion Z-test was performed in each cluster for E8.5 datasets (MLP: *P* = *0.655,* aSHF+SoM: *P* = *0.1829,* BrM: *P* = *0.477,* pSHF: *P* = *0.416*, CM: *P* = *0.790*). A two proportion Z-test in each cluster for E9.5 datasets (MLP: *P* < *2.2e*^−*16*^, aSHF+SoM: *P* = *0.7021,* BrM: *P* < *2.2e*^−*16*^, pSHF: *P* = *9.82e*^−*3*^, CM: *P* = *1.68e*^−*8*^). **J.** Single-cell embedding graph of the CPM lineages from *Tbx1^Cre^;Tbx1* Ctrl and *Tbx1^Cre^;Tbx1* cKO at E8.5 and E9.5 datasets colored by expression level. The color spectrum from blue to yellow indicates expression levels from low to high. Grey indicates no expression.

As in the integrated analysis of the four time point datasets shown in Figs. 1 and 2, the single timepoint dataset of the *Mesp1*+ cell lineage at E9.5 had similar cell populations (Fig. 4c, f, Extended Data Fig. 3a-e). Integrated and clustering analysis of the E9.5 datasets resolved CPM subpopulations better as compared to Figs. 1 and 2, as it could also identify the somatic mesoderm (SoM; Figure 4f, Extended Data Fig. 3d). Somatic mesoderm in embryos gives rise to the ventral pericardial wall (expressing *Msx1, Msx2*, *Epha3*, but not *Nkx2-5*^34^).

Comparison of the *Mesp1* and *Tbx1* lineages revealed that the MLPs are a derivative of *Tbx1* expressing cells within the *Mesp1* lineage (Figure 4c, g). There are some differences between the two *Cre* lines with respect to other lineages. The *Mesp1* lineage contributes more broadly to the embryonic mesoderm, while *Tbx1* is expressed in pharyngeal endoderm and distal pharyngeal ectoderm^35^ in addition to the CPM^36^. In the *Tbx1* cell lineage dataset (Fig. 4g, j, Extended Data Fig. 3f-j), we identified CPM, endoderm and ectoderm cells based upon marker genes and differential gene expression patterns (Supplementary Table 3). Although *Tbx1* is strongly expressed in the CPM, it is not expressed in the heart, neither in the FHF nor the caudal and medial pSHF at the timepoints analyzed^23,37,38^. Therefore, compared to the *Mesp1*+ lineage, there are fewer cells in pSHF and CM clusters in the *Tbx1* cell lineage due to the more limited expression domain of *Tbx1* (Fig. 4e, i, Ctrl).

Both *Mesp1^Cre^* and *Tbx1^Cre^* mediated *Tbx1* conditional null embryos, referred together as *Tbx1* cKO embryos, at E9.5, exhibited similar phenotypes including hypoplasia of the caudal pharyngeal apparatus comprising pharyngeal arches 3-6^36^. After cell clustering of the integrated datasets from control and *Tbx1* cKO embryos (*Mesp1* and *Tbx1* lineages), we analyzed the proportion of cell numbers in each cluster. We found a significant increase in the proportion of MLPs in *Tbx1* cKO embryos at E9.5 (Fig. 4d, e, h, i), but not in *Tbx1^Cre^;Tbx1* cKO embryos at E8.5 (Fig. 4h, i). In *Tbx1* cKO embryos at E9.5, the BrM and CM/OFT populations were significantly smaller than in controls. We noted that the pSHF population was increased in proportion in the *Mesp1^Cre^;Tbx1* cKO dataset but not in the *Tbx1^Cre^;Tbx1* cKO dataset. These results suggest that there are changes in the proportion of particular CPM and BrM cell populations upon inactivation of *Tbx1*. To investigate this further, we next tested whether there is also a change in gene expression within these CPM populations.

### *Tbx1* provides a balance of gene expression in the MLPs

To understand how *Tbx1* affects gene expression within the MLPs and derivative cell types, we analyzed differentially expressed genes (DEGs) with the scRNA-seq datasets. We analyzed DEGs in each cluster in control and *Tbx1* cKO embryos (*Mesp1^Cre^;Tbx1* Ctrl vs *Tbx1* cKO embryos at E9.5 [Supplementary Table 4] and *Tbx1^Cre^;Tbx1* Ctrl vs cKO embryos at E9.5 [Supplementary Table 5]). To focus on the most significant alterations and most biologically relevant genes, we examined only DEGs shared in both sets of embryos, which change in the same direction, with adjusted p-value < 0.05 and have an absolute log2 fold change > 0.25. In the MLPs, we identified 770 DEGs; 262 genes were decreased and 508 were increased in *Tbx1* cKO embryos (Fig. 5a, Supplementary Table 6). We also identified DEGs in MLP-derivative cells (Extended Data Fig. 4a-d). Among the 262 genes that were downregulated in MLPs (47 are MLP marker genes; Supplementary Table 1), a total of 104 genes were also downregulated in BrM (n=83), aSHF/SoM (n=41) and pSHF (n=20) cells in *Tbx1* cKO embryos (Fig. 5b, Extended Data Fig. 4j-l, Supplementary Table 7). Gene ontology (GO) analysis was used to identify enriched biological functions of the downregulated genes in *Tbx1* cKO embryos (Supplementary Table 8). Genes affected are involved in cell differentiation or cell signaling (e.g., *Nkx2-5, Tnnt2, Bmp7, Fgf10*; Fig. 5c). Taken together, these results suggest that *Tbx1* regulates expression of MLP genes required for lineage differentiation. *Tbx1* is also expressed in the BrM progenitor cells (C9; Fig. 1d, e, f). Some genes were specific for and downregulated only in the BrM population, including *Lhx2* and *Myf5* (Supplementary Table 7).

**Figure 5.**
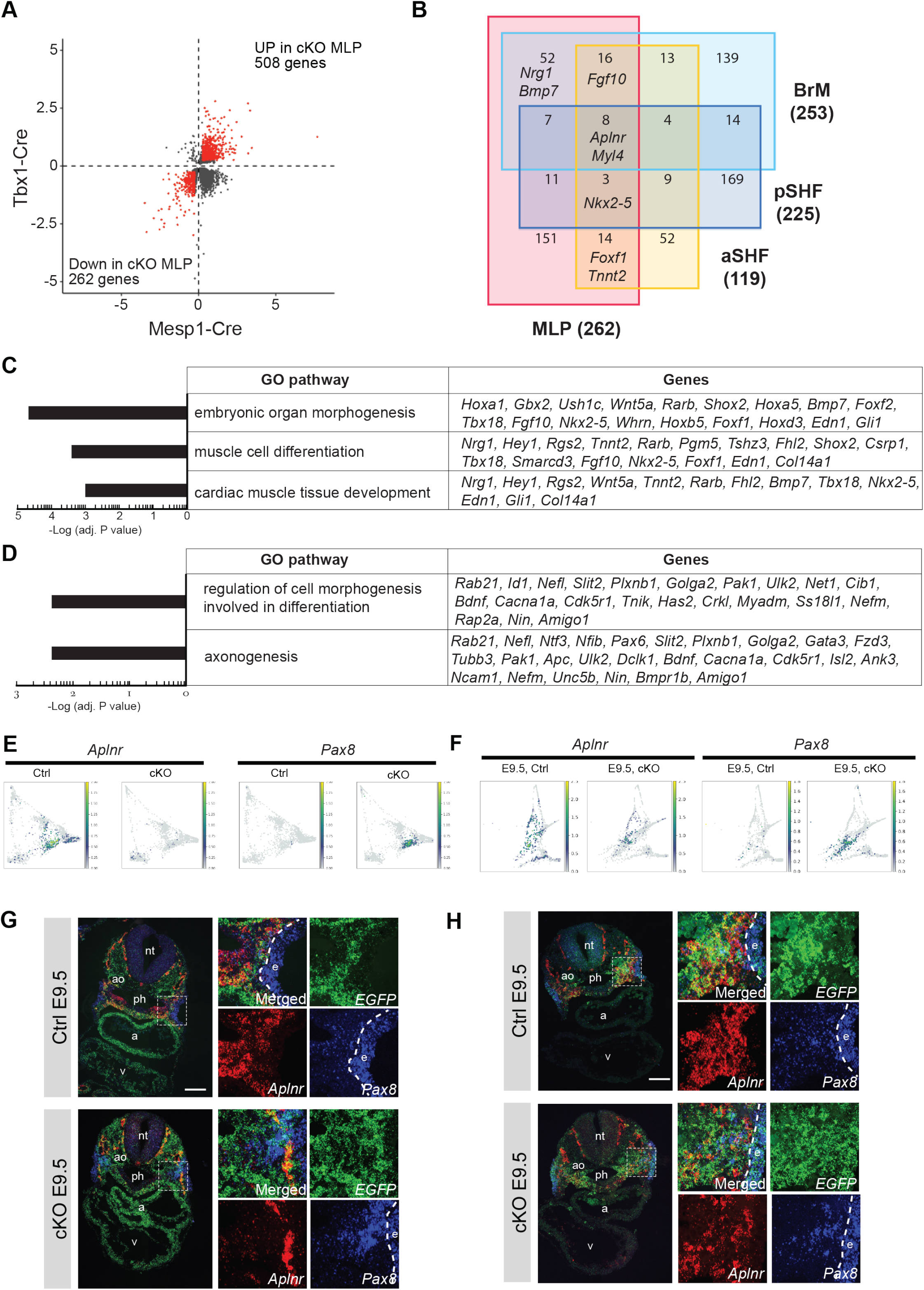
*Tbx1* is required in MLPs to promote lineage progression and restrict expression of ectopic genes. **A.** Comparison of differentially expressed genes (DEGs) from *Mesp1*^*Cre*/+^; *ROSA26-GFP*^*flox*/+^ (Ctrl) vs *Mesp1*^*Cre*/+^; *ROSA26-GFP*^*flox*/+^;*Tbx1*^*flox/flox*^ (cKO) embryos at E9.5 and *Tbx1*^*Cre*/+^; *ROSA26-GFP*^*flox*/+^ (Ctrl) vs *Tbx1*^*Cre/flox*^; *ROSA26-GFP*^*flox*/+^ (cKO) in MLPs. X-axis indicates log2-fold change of *Mesp1^Cre^* DEGs. Y-axis indicates log2-fold change of *Tbx1^Cre^* DEGs. Each dot indicates a gene, and red indicates common DEGs with |log2-fold change| > 0.25 in both comparisons. **B.** Venn diagram of downregulated genes in CPM clusters of *Tbx1* cKO embryos. The downregulated genes in MLP (Figure 4A), BrM, aSHF and pSHF (Extended Data Fig5B-E) populations were compared. The number in the diagram indicates the number of genes in each category. The full list is available in Supplemental Table 6-S10. **C** and **D.** Enriched biological processes in gene ontology (GO) and related genes found in downregulated (C) or upregulated (D) genes in MLPs of *Tbx1* cKO. X-axis indicates adjusted P values. **E.** Single-cell embedding graph of the CPM lineages from *Mesp1^Cre^;Tbx1* Ctrl and cKO at E9.5 datasets separated by genotype and colored by expression level. The color spectrum from blue to yellow indicates expression levels from low to high. Grey indicates no expression. **F.** Single-cell embedding graph of the CPM lineages from *Tbx1^Cre^;Tbx1* Ctrl and cKO at E9.5 datasets separated by genotype and colored by expression level. The color spectrum from blue to yellow indicates expression levels from low to high. Grey indicates no expression. **G.** RNAscope *in situ* hybridization with *EGFP*, *Aplnr* and *Pax8* mRNA probes on transverse section from *Mesp1*^*Cre*/+^;*ROSA26-GFP*^*flox*/+^;*Tbx1*^*flox*/+^ (Het, Ctrl) embryos (upper panel; n = 3) and *Mesp1*^*Cre*/+^;*ROSA26-GFP*^*flox*/+^;*Tbx1*^*flox/flox*^ (cKO) embryos (lower panel; n = 3) at E9.5. Green; *EGFP*, Red; *Aplnr*, Blue; *Pax8*. White dotted line indicates the position in higher magnification images shown in right. Upper left, merged image; Upper right, green channel (*EGFP*), Lower left, red channel (*Aplnr*); Lower right, blue channel (*Pax8*) Scale bar; 100 μm. **H.** RNAscope *in situ* hybridization with *EGFP*, *Aplnr* and *Pax8* mRNA probes on transverse section from *Tbx1*^*Cre*/+^;*ROSA26-GFP*^*flox*/+^ (Het, Ctrl) embryos (upper panel; n = 3) and *Tbx1*^*Cre/flox*^;*ROSA26-GFP*^*flox*/+^ (cKO) embryos (lower panel; n = 3) at E9.5. Green; *EGFP*, Red; *Aplnr*, Blue; *Pax8*. White dotted line indicates the position in higher magnification images shown in right. Upper left, merged image; Upper right, green channel (*EGFP*), Lower left, red channel (*Aplnr*); Lower right, blue channel (*Pax8*) Scale bar; 100 μm. Abbreviations: atrium (a), aorta (ao), ectoderm(e), neural tube (nt), pharynx (ph), ventricle (v).

Surprisingly, more genes were upregulated in *Tbx1* cKO MLPs (and other derivatives; Extended Data Fig. 5a-d; Supplementary Table 6) than downregulated (Fig. 5a, Supplementary Table 6). The enriched GO terms for the upregulated genes included axonogenesis, represented by key neuronal specific genes *Bdnf*^39^ or *Pax6*^40^ (Fig. 5d, Supplementary Table 8). Further, genes normally expressed in non-mesoderm cells were also expressed, such as *Pax8*^41^ (Supplementary Table 8). Therefore, inactivation of *Tbx1* results in both reduced expression of MLP genes for cell transitions and in increased/ectopic expression of non-mesodermal lineages, suggesting potential linage misspecification. We suggest that *Tbx1* provides a balance of specific gene expression required for MLP function.

To confirm expression changes from the scRNA-seq experiments when *Tbx1* is inactivated, we checked the expression pattern of *Aplnr* and *Pax8 in vivo* using RNAscope analysis. These genes are at the top fold change among DEGs decreased (*Aplnr)* and increased (*Pax8)* in the MLPs of *Tbx1* cKO embryos (Supplementary Table 4). Ectopic *Pax8* expression was observed in the MLPs of *Tbx1* cKO embryos (Fig. 5e, f, Extended Data Fig. 5a-d). In serial transverse sections of both *Tbx1* cKO and control embryos at E9.5, reduced levels of *Aplnr* expression in the MLPs in the posterior pharyngeal apparatus was observed, while *Pax8* was increased in the same region containing GFP+ cells (Fig. 5g, h, Extended Data Fig. 5e-h). We also examined DEGs and enriched GO terms in the other derivative clusters and found *Tbx1* also regulates expression in more differentiated cells (Extended Data Fig 4e-4i, Supplementary Table 9, 10). Overall, *Tbx1* provides a balance that promotes maturation but restricts ectopic expression of non-mesodermal genes in MLPs.

### TBX1 defines a gene regulatory network in the MLPs for cardiac and BrM formation

To better understand how TBX1 regulates the expression of genes in the CPM at the chromatin level, we performed ATAC-seq of control versus *Tbx1* mutant embryos (Fig. 6a, Extended Data Fig. 6a-c) and TBX1 ChIP-seq analysis at E9.5. We identified MLP enriched genes from DEGs for MLPs versus other CPM cell types. The ATAC-seq peaks from three replicates were separated to commonly accessible regions (CARs) or differentially accessible regions (DARs; FDR < 0.05, Fig. 5b, Extended Data Fig. 6d) between control and mutant samples. Among 5,872 DARs, 5,859 decreased and 13 increased in chromatin accessibility in *Tbx1* cKO embryos. To examine the CPM, we next performed ATAC-seq on *Mesp1*^*Cre*/+^ cells and used the peaks to exclude CARs and DARs that were found only in *Tbx1*^*Cre*/+^ and *Tbx1^Cre/flox^* samples to eliminate changes in non-mesoderm lineages (Fig. 6a, Extended Data Fig. 6a-c). The remaining CARs and DARs are thus referred to as “CARs-Mesp1” and “DARs-Mesp1” (Fig. 6c, d). A total of 81.4% of CARs-Mesp1 were in promoter regions, while 35.8% of DARs-Mesp1 were in promoter regions and 34.9% were in distal intergenic regions (Extended Data Fig. 6e, f), suggesting that *Tbx1* inactivation has a large effect on regulation of genes at a distance from their promoters.

**Figure 6.**
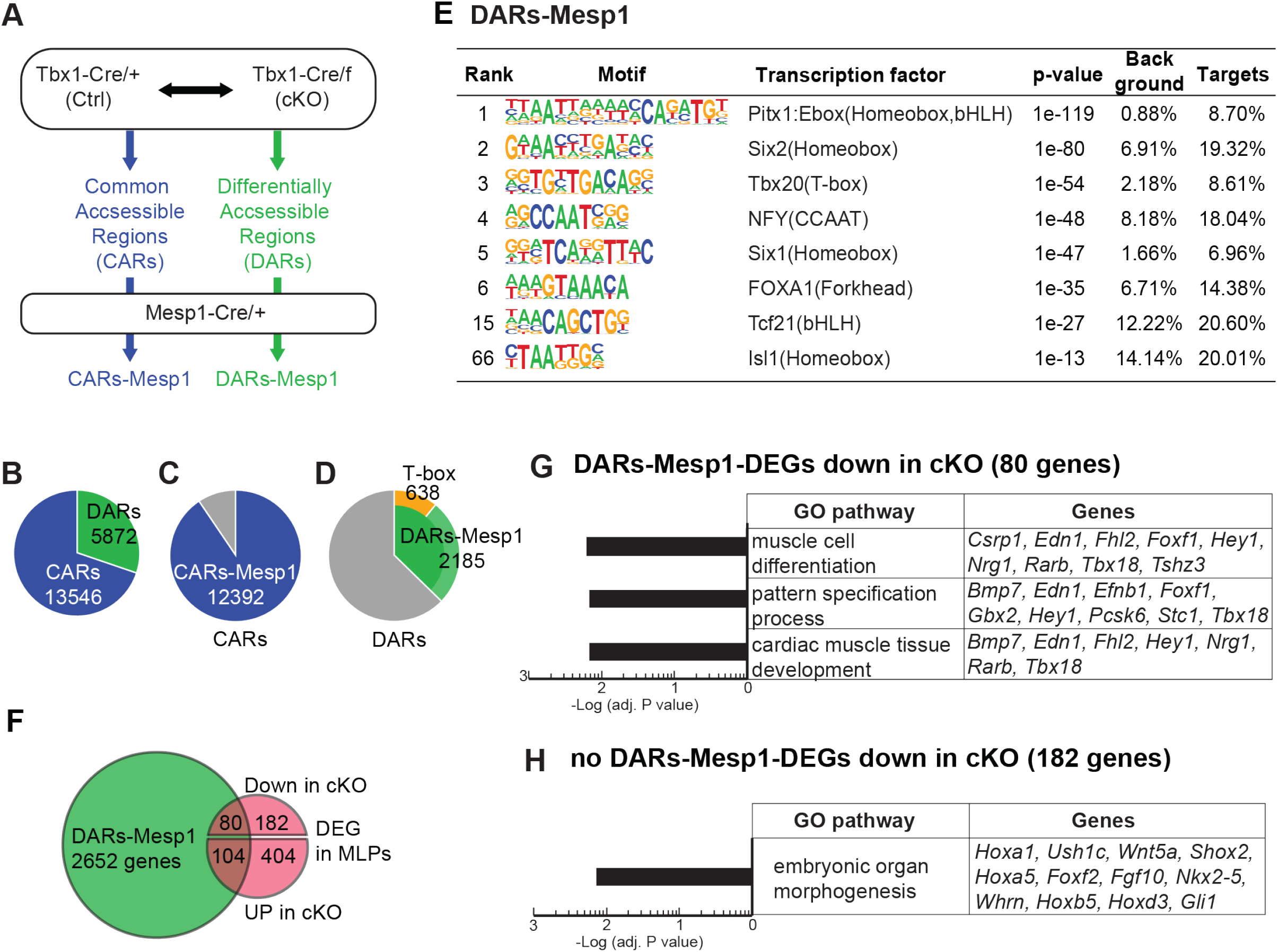
*Tbx1* affects gene expression and chromatin accessibility of MLPs. **A.** Scheme of ATAC-seq data analysis. **B.** Pie chart of ATAC-seq peaks of GFP+ cells from *Tbx1*^*Cre*/+^; *ROSA26-GFP*^*flox*/+^(Ctrl) and *Tbx1*^*Cre/flox*^; *ROSA26-GFP*^*flox*/+^ (cKO) embryos at E9.5 (n = 3). A total of 13,546 peaks, defined as CARs, commonly accessible regions, were found in both *Tbx1^Cre^ Tbx1* Ctrl and cKO datasets. A total of 5,872 peaks, defined as DARs, differentially accessible regions, in which peak intensity was changed (FDR ≤ 0.05) between *Tbx1^Cre^ Tbx1* Ctrl and cKO datasets. **C.** Pie chart showing the intersection of CARs with ATAC-seq peaks of GFP+ cells from *Mesp1*^*Cre*/+^; *ROSA26-GFP*^*flox*/+^ embryos at E9.5 (n = 3). Among 13,546 CARs, a total of 12,392 peaks were also found to open in *Mesp1^Cre^* lineages and thus referred as CARs-Mesp1. **D.** Pie chart showing the intersection of DARs with ATAC-seq peaks of GFP+ cells from *Mesp1*^*Cre*/+^;*ROSA26-GFP*^*flox*/+^ embryos at E9.5 (n = 3). Among 5,872 DARs, 2,185 were also found to open in *Mesp1^Cre^* lineages and thus referred as DARs-Mesp1. Of the DARs-Mesp1 regions, 638 had a T-box motif. **E.** The motifs found in DARs-Mesp1 regions by Homer (Hypergeometric Optimization of Motif Enrichment). **F.** Venn diagram of genes with ATAC-seq DARs-Mesp1 regions and DEGs in MLPs from scRNA-seq. Of the 770 DEGs in the MLPs, 80 genes showed decreased and 104 genes showed increased chromatin accessibilities in *Tbx1* cKO embryos. **G.** Enriched biological processes for the 80 genes with both DARs-Mesp1 and reduced expressed in *Tbx1* cKO MLPs. X-axis indicates adjusted P values. **H.** Enriched biological processes for the 182 genes with reduced expression in *Tbx1* cKO but without DARs-Mesp1. X-axis indicates adjusted P values.

We then focused upon MLP genes within the CPM. A total of 80 of the 262 downregulated genes in the MLPs in *Tbx1* cKO embryos were associated with DARs-Mesp1 (Fig. 6f). In DARs-Mesp1, several transcription factors binding motifs related to heart or BrM differentiation including the T-box motif were identified, and many of these are likely MLP relevant transcription factors (Fig. 6e). We also used GREAT^42^ to identify genes that harbored DARs-Mesp1 and their enriched functions. GO analysis of the 80 genes indicated that these genes were involved in skeletal muscle cell differentiation, pattern specification process and cardiac muscle tissue development, important for the function of *Tbx1* (Fig. 6g). On the other hand, those without DARs-Mesp1 were not specific to the CPM populations (Fig. 6h). Despite the fact that more genes are upregulated when *Tbx1* is inactivated, they contained only 13 DARs that increased in accessibility, and none were associated with the DARs-Mesp1. Therefore, *Tbx1* might indirectly affect these genes in the MLPs that were not detected by ATAC-seq analysis.

To determine which genes with DARs could be direct target genes of TBX1, we performed ChIP-seq with an Avi-tagged *Tbx1* mouse line that we generated, and created double homozygous mice harboring the biotin ligase (BirA) gene (*Tbx1^3’-Avi^;BirA*, Extended Data Fig. 7a). We considered peaks found in at least two of the three replicates as high confident TBX1 binding sites (Extended Data Fig. 7b, c). Out of a total of 255 peaks, 176 peaks had a T-box motif (Fig. 7a, Extended Data Fig. 7d, e), supporting TBX1 occupancy; 104 peaks (41%) in the ChIP-seq data overlapped with DARs (Fig. 7a, c, Extended Data Fig. 7f). Comparing *Tbx1*^*Cre*/+^ and *Tbx1^Cre/flox^* ATAC-seq data in 255 ChIP-seq peak regions, we found that these regions were mostly open and accessible in *Tbx1*^*Cre*/+^ embryos and closed in *Tbx1^Cre/flox^* embryos (Fig. 7c, Extended Data Fig. 7f). Of the 255 ChIP-seq peaks, 151 (59%) did not show significant chromatin accessibility changes (i.e., overlapping with DARs), which include some TBX1 binding sites that were located in closed regions that did not change in accessibility in the mutant data (Fig. 7c), suggesting a diverse role of TBX1 in promoting chromatin remodeling.

**Figure 7.**
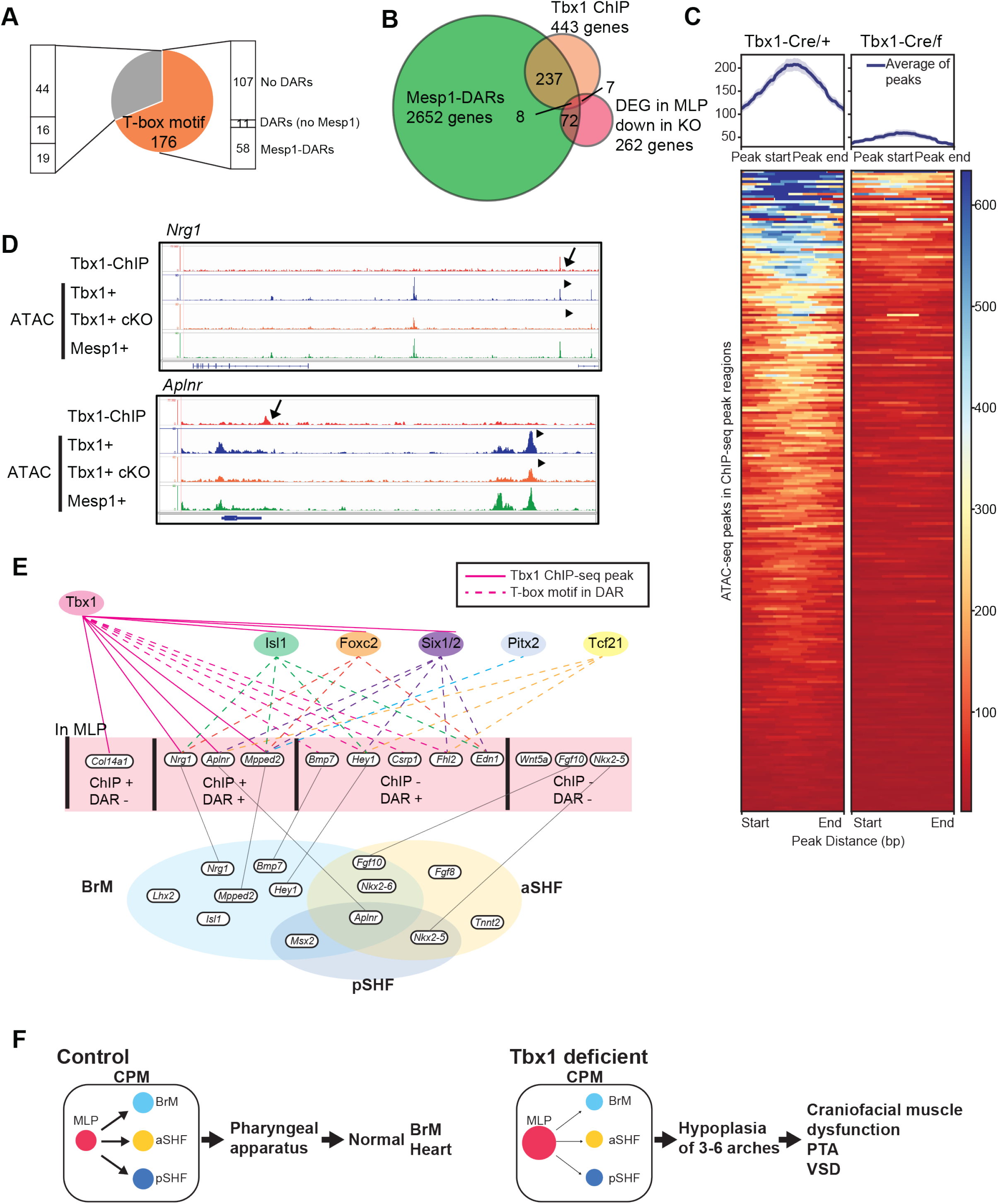
Open chromatin status in TBX1 binding regions *in vivo* were reduced in *Tbx1* cKO embryos. **A.** Pie chart of TBX1 ChIP-seq peaks from three biological replicates. Among 255 ChIP-seq peaks, 176 had a T-box motif. The bar chart indicates peak intersection with ATAC-seq data. **B.** Venn diagram for overlap of genes with ChIP-seq peaks or ATAC-seq DARs-Mesp1 regions or downregulated expression in *Tbx1* cKO MLPs. **C.** Average chromatin accessibility at ChIP-seq peaks. Left, ATAC-seq data from *Tbx1* Het embryos; Right, ATAC-seq data from *Tbx1* cKO embryos. Top, average ATAC-seq signal in TBX1 ChIP-seq peak regions; Bottom, heatmap of ATAC-seq read densities in TBX1 ChIP-seq peaks. Statistical significance (adj. P < 0.0009) of ChIP-seq peaks on ATAC-seq signal has been tested using enrichPeakOverlap present in ChIPseeker. **D.** Genome browser snapshots including results from TBX1 ChIP-seq, ATAC-seq of *Tbx1* Het embryos, ATAC-seq of *Tbx1* cKO embryos, ATAC-seq from Mesp1+ lineages, at *Nrg1* (upper), *Aplnr* (lower) loci. Arrow indicates TBX1 ChIP-seq peak. Arrowhead indicates DARs. **E.** A cartoon of the TBX1 regulation network in the MLPs and derivatives. Pink lines indicate TBX1 direct target genes from *in vivo* ChIP-seq. Dotted lines indicate genes with the TBX1 motif harboring DARs-Mesp1 from ATAC-seq analysis. The transcription factor genes indicated in different colors (*Isl1, Foxc2, Six1/2, Pitx2, Tcf21*) are representative family members sharing the same binding motif enriched in DARs-Mesp1. The genes were identified in DEGs of MLP, BrM, aSHF and pSHF populations from scRNA-seq analysis. Grey line indicates overlapping genes found in MLPs and other differentiated branches. **F.** A working hypothesis of MLP function with respect to *Tbx1*. In *Tbx1* conditional null embryos, the MLPs fail to properly transition to more differentiated CPM states. This leads to an accumulation of MLPs and reduction of more differentiated cell types (BrM, aSHF, pSHF) leading to fewer CMs. Fewer BrM cells result in intermittent failure of these muscles to form, while fewer CMs lead to a shortened cardiac outflow tract that results in a persistent truncus arteriosus (PTA) with an obligate ventricular septal defect (VSD) in the heart. Further, the deficient CPM has non-autonomous effects on the neighboring neural crest cells and epithelial cells, resulting in hypoplasia of the distal pharyngeal apparatus with failed segmentation to pharyngeal arches 3, 4 and 6 (there is no pharyngeal arch 5 in mammals).

We intersected the genes with DARs-Mesp1 and the 262 genes reduced in the MLPs in *Tbx1* KO embryos to identify MLP specific gene regulation. When we further intersected the ATAC-seq and TBX1 ChIP-seq, we found eight genes (Fig. 7b), including *Aplnr* and *Nrg1*, which are MLP enriched genes that are TBX1 direct transcriptional targets (Fig. 7d). In the *Nrg1* locus, the TBX1 binding regions were closed in *Tbx1* cKO embryos, while in the *Aplnr* gene region, the TBX1 binding site was not in a DAR (Fig 7d), indicating that multiple mechanisms of regulation occur.

Taking the results from the three types of functional genomic data in this report, we can generate a putative gene regulatory network for TBX1 function in the MLPs as summarized in Figure 7e. Here, we distinguish four categories of genes regulated by TBX1: 1) Direct target genes with or 2) without chromatin changes, and indirect target genes 3) with chromatin changes, that contain transcription factor binding sites and 4) without chromatin changes. Note that some DEGs changed expression in more than one cell type or population as indicated (gray lines). Overall, we suggest that TBX1 with *Isl1, Fox, Six, Pitx* and E-box proteins such as *Tcf21,* (Fig. 6e, 7e), act together to regulate progression of MLPs to more differentiated states in the CPM.

## Discussion

Single cell RNA-seq is uncovering MLPs in different developmental contexts, these cells serving as progenitor states in organogenesis. In this report, we uncovered the cellular and molecular processes by which a specific MLP population functions to both maintain a progenitor state and allocate cells to derivative cell types. We discovered that MLPs are localized bilaterally within the posterior nascent mesoderm of the pharyngeal apparatus. The MLPs are needed for deployment of progenitor cells to derivative tissues during the time when individual arches are forming in a rostral to caudal manner, thereby elongating the pharyngeal apparatus. We focused on function of *Tbx1*, the gene for 22q11.2DS, which marks the MLP lineage in the CPM. In Figure 7F, we show a model for *Tbx1* function. Inactivation of *Tbx1* results in dysregulation of gene expression in MLPs, and reduced progression to more mature states. In turn, this causes an accumulation of MLPs and reduction of more differentiated CPM cells, the latter resulting in fewer cardiomyocytes leading to a shortened outflow tract. To better understand TBX1 function we performed ATAC-seq and TBX1 ChIP-seq, and we defined a gene regulatory network in the CPM in which TBX1 plays a key role, thereby helping to explain the cause of the cardiac and BrM phenotype in 22q11.2DS patients.

### Dynamic cell allocation of MLPs to the heart and BrMs

An example of how MLPs function in the mammalian mesoderm during organogenesis is observed during somite formation in the tail of vertebrate embryos where the neuromesodermal progenitors provides a continuous source of cells to form segmented somites and neural tube ^43^. In this report, the MLPs are required during segmentation of the pharyngeal apparatus to individual arches. The presomitic mesoderm forms after gastrulation and this is when the progenitors of MLPs form in the embryo. Subpopulations of the CPM are also specified during gastrulation ^8^. At gastrulation, multilineage primed cells expressing genes of pluripotent markers, aSHF and pSHF progenitor genes can be identified^8^. The MLPs in this report are part of the CPM, and do not express pluripotency markers, but need to continue to function as a constant source of progenitors. The segmentation of the pharyngeal apparatus is an extremely dynamic process in which surrounding signals are constantly changing. The MLPs, located posterior and lateral to the developing heart allocate cells during this dynamic process thereby continuously providing cells to the cardiac poles and for formation of the BrMs, in the ever-changing local environment.

One question is how the MLPs can provide cells to the poles of the heart, while located behind (dorsal) to the heart tube. The cells deployed to the cardiac poles first arrive at the dorsal pericardial wall (DPW), just behind the heart tube ^44^. The cells of the CPM must undergo mesenchymal-epithelial transition to form the DPW^44^. The mesoderm cells lateral and behind the DPW are incorporated to the DPW and then to the poles of the heart^45^. The deployment of these adjacent mesoderm cells to the DPW provides a pushing force as the epithelial transitioned cells move to the poles of the heart^45,46^.

Our *in vivo* localization of MLP markers in the mouse embryo suggest that the MLPs are the dorsal population of mesoderm that are needed to allocate cells to the DPW. In addition to deploying cells to the DPW, the MLPs must also deploy cells to the BrMs. The BrMs form segmentally in a rostral to caudal manner, in which first the muscles of mastication form from the first arch and the other muscles of the face and neck form thereafter from the caudal arches. The BrMs express myogenic regulatory transcription factors including *Tcf21*, *Msc*, *Myf5* and later *Myod1*^47^. In addition, transcription factor genes such as *Isl1*^11^, *Pitx2^48^*, *Tbx1*^19^, *Lhx2*^24^ and *Ebf* genes (*Ciona*^49^), among others, are expressed in the CPM and required for BrM formation. Loss of *Tbx1* results in intermittent failure of BrMs from forming^19^ and acts upstream of many of these important transcription factors^24^. This suggests that the *Tbx1* gene regulatory network is central in MLPs and, independently to BrM specification.

### Molecular mechanism for *Tbx1* function

In this report, we identify a novel function of *Tbx1*, in a newly recognized CPM subpopulation termed MLPs. We found that *Tbx1* is required for the MLPs to progress, particularly for the aSHF, CM and BrM fates. Therefore, one of the main functions of the MLPs is to allocate CPM cells to more differentiated states. Second, we found that *Tbx1* contribution is for the aSHF, CM and BrM cells and less so for the *Tbx5* expressing pSHF cells^25^, consistent with the expression pattern of *Tbx1*. To understand how the MLPs transition and with respect to *Tbx1*, we defined a gene regulatory network which provides a molecular mechanism that includes new genes for CPM function during the acquisition of aSHF, CM and BrM fates. By performing multi-omic studies, we identified *Aplnr* and *Nrg1* among the genes enriched in expression in MLPs. *Aplnr* encodes the APJ/Apelin G-protein coupled receptor that binds Apelin or Elabela/Toddler peptide ligands that have many embryonic and adult functions^52^. In zebrafish, knockdown of *Aplnr* disrupts normal migration of cells during gastrulation including that of cardiac progenitors resulting in severe defects^53,54^. Unexpectedly, their role in early embryogenesis is not recapitulated fully in mouse models, implicating perhaps functional redundancy with other G-protein coupled receptors or ligands^55^. *Aplnr* is expressed in the CPM, and from stem cell studies, has a role in cardiomyocyte development^29^. *Nrg1* is also of particular interest. In contrast to *Aplnr, Nrg1* is not expressed in the DPW. *Nrg1* encodes an EGF family ligand that binds to ErbB receptor tyrosine kinases and has multiple roles in cardiac development and function^30,56^. Interestingly, both *Aplnr* and *Nrg1* are direct target genes of TBX1 based on our ATAC-seq and ChIP-seq results, suggesting that these genes are mediators of TBX1 function in MLPs.

Some of the genes that were differentially expressed and differentially accessible in *Tbx1* conditional null embryos include *Isl1*^57,58^, *Foxc2*^59,60^ *Six1* or *Six2*^61^, *Pitx2*^48,62^ and *Tcf21*^24^. *Isl1*, *Foxc2* and *Six2* may be direct transcriptional target genes of TBX1 based upon ChIP-seq analysis. Some known downstream genes of *Tbx1* were not identified in the multi-omic data, such as *Wnt5a*^63^, *Fgf10*^64,65^ and *Nkx2-5*, possibly due to low transcript abundance. A subset of the genes with differentially accessible regions are altered in MLPs as well as derived cell types. This is particularly true for the BrM progenitor cells, where *Tbx1* is actively expressed. Note that our finding of chromatin status changes in *Tbx1* mutant embryo is different as compared to analysis of cell culture^66^, suggesting differences between cell culture versus the *in vivo* context or differentiation stages.

## Conclusions

In this study, we identified MLPs as a continuous but evolving source of CPM cells that is maintained during development of the pharyngeal apparatus. *Tbx1* marks the MLP cell population in embryogenesis. Further, *Tbx1* is required in MLPs to promote their maturation to more differentiated states by direct and indirect regulation of chromatin accessibility and transcriptional regulation.

## Online Methods

### Mice

All experiments using mice were carried out according to regulatory standards defined by the National Institutes of Health and the Institute for Animal Studies, Albert Einstein College of Medicine (https://www.einstein.yu.edu/administration/animal-studies/), IACUC protocol is #0000-1034.

The following mouse mutant alleles used in this study have been previously described: *Mesp1^Cre^* ^7^, *Tbx1^Cre^* ^67^, *Tbx1^flox/flox^* ^68^, *ROSA26-GFP^flox/flox^* (RCE: loxP)^69^. For lineage tracing in *Tbx1* conditional knockout embryos, *ROSA26-GFP^flox/flox^;Tbx1^flox/flox^* and *ROSA26-GFP*^*flox/flox*^;*Tbx1*^*flox*/+^ mice were generated by inter-crossing *Tbx1^flox/flox^* and *ROSA26-GFP^flox/flox^* mice. To generate *Mesp1*^*Cre*/+^; *ROSA26-GFP*^*flox*/+^ embryos, *Mesp1*^*Cre*/+^ male mice were crossed with *ROSA26-GFP^flox/flox^* female mice. To generate *Mesp1^Cre/+^; ROSA26-GFP^flox/+^;Tbx1^flox/flox^* mutant embryos, *Mesp1*^*Cre*/+^ transgenic mice were crossed with *Tbx1^flox/flox^* mice to obtain *Mesp1*^*Cre*/+^;*Tbx1*^*flox*/+^ male mice, then crossed with *ROSA26-GFP^flox/flox^;Tbx1^flox/flox^* female mice. The littermates with *Mesp1*^*Cre*/+^; *ROSA26-GFP*^*flox*/+^;*Tbx1*^*flox*/+^ were used as Ctrl in RNAscope experiment. To generate *Tbx1^Cre/+^; ROSA26-GFP^flox/+^* and *Tbx1^Cre/flox^;ROSA26-GFP^flox/+^* embryos for scRNA-seq and ATAC-seq, *ROSA26-GFP^flox/flox^* female mice and *ROSA26-GFP^flox/flox^;Tbx1^flox/flox^* female mice were crossed with *Tbx1*^*Cre*/+^ male mice, respectively. For RNAscope analysis, littermates were used from a cross of *Tbx1*^*Cre*/+^ male mice and *ROSA26-GFP*^*flox/flox*^;*Tbx1*^*flox*/+^; female mice. All the mice are maintained on the SwissWebster genetic background. The PCR strategies for mouse genotyping have been described in the original papers and are available upon request.

To generate *Tbx1^3’-Avi^* mice, Avidin (Avi) was inserted using CRISPR/Cas9 genomic engineering in the Gene Modification Facility of Albert Einstein College of Medicine. A guide RNA (gRNA) targeting to the last exon of *Tbx1* (5’-gcgcgcgggcgcactatctgggg-3’) was designed by Guide Design Resources (http://crispr.mit.edu/) and generated by *in vitro* transcription^69^. Cas9 mRNA was purchased from SBI. *Tbx1^3’-Avi^* homology directed repair (HDR_ vector containing the 60 nt homologous arms (5’-ggcggccgcgccgcccggtgcctacgactactgccccagaGGTGGAAGTggcctgaacgacatcttcgaggctcagaaaatcgaatggcacgaatagtgcgcccgcgcgccgaccccgagggccatccaaggacgcgctccc-3’) at each side surrounding the Avi tag was synthesized chemically from IDT. Super-ovulated female C57BL6 mice (3–4 weeks old) were crossed with C57BL/6 males, and fertilized embryos were collected from oviducts. The gRNA, Cas9 mRNA and *Tbx1^3’-Avi^* HRD vectors were mixed and microinjected into the cytoplasm of fertilized eggs. The injected zygotes were transferred into pseudo pregnant CD1 females, and the resulting pups were obtained. For genotyping, 311 bp (wildtype) and 365 bp (*Tbx1^3’-Avi^*) bands were observed with *Tbx1^3’-Avi^* F4: 5’- gcagccaacgtgtactcgtc-3’ and *Tbx1^3’-Avi^* R2: 5’-gccggtgcagtatctacagt-3’ primer pair. After several backcrosses with Swiss Webster mice, *Tbx1^3’-Avi^* mice were crossed with *FVB;129P2-Gt(ROSA)26Sor^tm1.1(BirA)Mejr^/J* (*BirA*) mice from the Jackson laboratory to obtain *Tbx1^3’-Avi^;BirA* double homozygous mice.

The embryos at E8.0-E10.5 determined by standard somite counts, were used for all the experiments. Because of the difficulty to distinguish the sex at those time points, we used both male and female embryos for all the experiments.

### scRNA-seq

The embryos at E8.0-10.5 were isolated and GFP positive embryos were selected under a SteREO Discovery.V12 microscope (Carl Zeiss, Jena, Germany) in ice-cold DPBS with Ca^2+^ and Mg^2+^ (GIBCO, Cat# 14040-133). The rostral half of the embryos were collected at E8.0, E8.25 and E8.5. The pharyngeal apparatus with heart was collected at E9.5 and E10.5, as shown in Fig. 1A. The microdissected tissues were pooled in DMEM (GIBCO, Cat# 11885-084) until all the dissections were completed. Following centrifugation (4 °C, 100 x g, 5’, minutes) and removal of DMEM, tissues were incubated with 0.25% Trypsin-EDTA (GIBCO, Cat# 25200-056) with Dnase I (50U/ml) (Millipore, Cat# 260913-10MU), 10’ at RT. Then FBS (heat-inactivated, ATCC, Cat# 30-2021) was added to stop the reaction. After centrifugation (4 °C, 300 x g, 5’), the cells were resuspended in PBS w/o Ca^2+^ and Mg^2+^ (Corning, Cat# 21-031-cv) with 10% FBS and passed through the 100 μm cell strainer. DAPI (Thermo Fisher Scientific, Cat# D3571) was added before cell sorting. The GFP+DAPI-cells were sorted with the BD FACSAria II system (Becton, Dickinson and Company, Franklin Lakes, NJ). The sorted GFP positive cells were centrifuged (4 °C, 300 x g, 5’), and resuspended in 50 μl PBS w/o Ca^2+^ and Mg^2+^ with 10% FBS. After measuring cell number and cell viability, the cells were loaded in a 10x Chromium instrument (10x Genomics, Pleasanton, CA) using Chromium Single Cell 3’ Library & Gel Bead Kit v2 (10x Genomics, Cat# PN-120237) or Chromium Next GEM Single Cell 3’ GEM, Library & Gel Bead Kit v3.1 (10x Genomics, Cat# PN-1000121) according to the manufacturer’s instructions (Genomics Core Facility). The concentrations of the libraries were measured with Qubit®2.0 Fluorometer (Thermo Fisher Scientific, Waltham, MA) using Qubit dsDNA HS Assay Kit (Thermo Fisher Scientific, Cat# Q32851). The details of each sample are provided in Table 1.

### ATAC-seq

The ATAC-seq method has been previously described^70^. The embryos at E9.5 were isolated from a euthanized mother. GFP positive embryos were selected under a SteREO Discovery.V12 microscope in ice-cold DPBS with Ca^2+^ and Mg^2+^ (GIBCO, Cat# 14040-133). The pharyngeal apparatus without the heart was collected and pooled in DMEM (GIBCO, Cat# 11885-084) until all the dissections were completed. Following centrifugation (4 °C, 100 x g, 5’) and removal of DMEM, tissues were incubated with 0.25% Trypsin-EDTA (GIBCO, Cat# 25200-056) with Dnase I (50U/ml) (Millipore, Cat# 260913-10MU), 10’ at RT. Then FBS (ATCC, Cat# 30-2021) was added to stop the reaction. After centrifugation (4 °C, 300 x g, 5’), the cells were resuspended in PBS w/o Ca^2+^ and Mg^2+^ (Corning, Cat# 21-031-cv) with 3% FBS and passed through the 100 μm cell strainer. The DAPI (Thermo Fisher Scientific, Cat# D3571) was added before the sorting. The GFP+DAPI-cells were sorted with the BD FACSAria II system (Becton, Dickinson and Company, Franklin Lakes, NJ). The sorted cells were centrifuged (4 °C, 800 x g, 5’), and washed with PBS w/o Ca^2+^ and Mg^2+^. After centrifugation (4 °C, 800 x g, 5’), cells were resuspended with ice-cold Lysis buffer which contained 10 mM Tris-HCl pH 7.4 (Thermo Fisher Scientific, Cat# BP152-1), 10 mM NaCl (Thermo Fisher Scientific, Cat# AM9760G), 3 mM MgCl2 (MilliporeSigma, Cat# M9272) and 0.1% Igepal CA-630 (MilliporeSigma, Cat# I8896). Following centrifugation (4 °C, 1000 x g, 5’), nuclear pellets were incubated with Tn5 transpose from the Nextera DNA Sample Preparation Kit (Illumina, Cat# FC-121-1030) for 37°C, 30’. Transposed DNA was purified using the MinElute PCR purification kit (QIAGEN, Cat# 28004), according to manufacturer’s instructions. The transposed DNA was amplified with NEB Net High Fidelity 2x PCR master mix, PPC, and two index primers from the Nextera Index Kit (Illumina, Cat# FC-121-1011). PCR conditions for amplification is 1 cycle 5’ 72 °C, 30” (seconds) 98 °C, 12 cycle 10”, 98 °C, 30”, 63 °C, 1’, 72 °C. PCR products were purified using the MinElute PCR purification kit. DNA concentration was measured with a Qubit®2.0 Fluorometer using Qubit dsDNA HS Assay Kit. DNA qualities were analyzed with a Bioanalyzer (Agilent Technologies, Santa Clara, CA).

### ChIP-seq

Whole embryos from *Tbx1^3’-Avi^;BirA* double homozygous embryos or *BirA* homozygous embryos (Ctrl) at E9.5 were collected and minced in ice-cold PBS. After centrifugation (4 °C, 200 x g, 5’), tissues were cross-linked with 1% formaldehyde (Thermo Fisher Scientific, Cat# 28906), 30’ at RT. A total of 2.5 M glycine (MilliporeSigma, Cat# G8898) was added at a final concentration 0.125 M to stop the reaction. After the tissues were washed with PBS, and centrifuged (4 °C, 200 x g, 5’), the pellets were frozen in dry ice and stored at −80 °C. We used 20 embryos for one sample. The frozen tissues were homogenized by BeadBug Microtube Homogenizer (Benchmark Scientific, Edison, NJ) in Lysis buffer, which contained 50 mM Hepes pH 7.5 (MilliporeSigma, Cat# H0887), 140 mM NaCl, 1 mM EDTA (MilliporeSigma, Cat# E7889), 10% Glycerol (MilliporeSigma, Cat# 4750-OP), 0.5% IGEPAL® CA-630 (MilliporeSigma, Cat# I8896), 0.25% TritonX-100 (MilliporeSigma, Cat# X100) and Protease Inhibitor cocktail (MilliporeSigma, Cat# P8340). Samples were incubated on ice for 10’. After centrifugation (4 °C, 2000 x g, 5’), extracted nuclei were washed in Wash buffer containing 10 mM Tris-HCl pH 8.0 (MilliporeSigma, Cat# T2694), 200 mM NaCl, 1mM EDTA pH 8.0, 0.5 mM EGTA pH 8.0 (MilliporeSigma, Cat# E3889) and Protease Inhibitor cocktail. Then the nuclei were resuspended in Shearing buffer contained 10 mM Tris-HCl pH 8.0, 1mM EDTA pH 8.0, 0.1% SDS (BIO-RAD, Cat# 1610418) and Protease Inhibitor cocktail. Resuspended nuclei were sonicated with a S2 Focused-ultrasonicator (Covaris, Inc., Woburn, MA). After sonication, 10% TritonX-100 and 5M NaCl were added to sheared chromatin in a final concentration of 1% TritonX-100 and 150 mM NaCl. Dynabeads™ MyOne™ Streptavidin T1 (Thermo Fisher Scientific, Cat# 65602) were blocked with SEA BLOCK Blocking Buffer (Thermo Fisher Scientific, Cat# 37527) and sheared chromatin was blocked with Dynabeads™ Protein A for immunoprecipitation (Thermo Fisher Scientific, Cat# 10002D) and Dynabeads™ Protein G for immunoprecipitation (Thermo Fisher Scientific, Cat# 10004D) at 4 °C, 1 hour on a rotator.

Following washing with Shearing buffer, blocked streptavidin magnetic beads were added to the sheared chromatin, then incubated 4°C overnight on a rotator. Streptavidin magnetic beads were washed with Low salt wash buffer (20 mM Tris-HCl pH 8.0, 45 mM NaCl, 0.1% SDS, 1% Triton X-100, 2mM EDTA), High salt wash buffer (20 mM Tris-HCl pH 8.0, 150 mM NaCl, 0.1% SDS, 1% Triton X-100, 2mM EDTA), LiCl wash buffer containing 10 mM Tris-HCl pH 8.0, 0.25M LiCl (MilliporeSigma, Cat# L7026), 1% IGEPAL® CA-630,1% sodium deocxycholate (MilliporeSigma, Cat# D6750), 1 mM EDTA. After a wash with TE buffer, DNA was eluted in Elution buffer (10 mM Tris-HCl pH 8.0, 1 mM EDTA, 1% SDS) with Proteinase K (Promega, Cat# V3021) at 65°C, 5 hours. Eluted DNA was purified with the MinElute PCR purification kit. After measuring the DNA concentration with the Qubit®2.0 Fluorometer using Qubit dsDNA HS Assay Kit, the libraries were prepared using Accel-NGS 2S Plus DNA Library Kit (Swift Bioscience, Cat# 21024) and 2S Set A Indexing Kit (Swift Bioscience, Cat# 26148) then the DNA concentration was measured with the Qubit®2.0 Fluorometer using Qubit dsDNA HS assay kit.

### Sequencing

The DNA libraries were sequenced using a Illumina HiSeq2500 system (at Einstein Epigenomics Core Facility), Illumina HiSeq4000 system (at Genewiz, South Plainfield, NJ) or NovaSeq6000 system (at Novogene, Sacramento, CA), with paired-end, 100 bp read length.

### RNAscope

After incubation of 4% PFA (Alfa Aesar, Cat# 43368) at 4 °C for 4 hours, embryos were dehydrated with 70%, 90% and 100% Ethanol (Thermo Fisher Scientific, Cat# A405P-4). Incubation with xylene (Thermo Fisher scientific, Cat# X3S-4) was done at RT, one hour twice followed by incubation of paraffin (Thermo Fisher Scientific, Cat# T555) at 65°C, one hour twice. Embryos were embedded and stored at 4°C.

Embryos were sectioned at 7 μm thickness and dried at RT. The sections were processed with RNAscope® Multiplex Fluorescent v2 (Advanced Cell Diagnostics, Cat# 323100), according to the manufacturer’s instructions. Briefly, sections were deparaffinized in xylene and dehydrated in graded ethanol, then incubated with hydrogen peroxide at RT, in 10’. The sections were incubated in boiled 1x Target retrieval reagent, for 15’. After rinsing with distilled water and 100% ethanol, the sections were dried at RT, for 5’. The sections were incubated with Protease Plus at RT, for 3’ then incubated with mixed RNAscope probes of C1, C2 and C3 channels (see Oligonucleotides section of the Table), at 40°C, overnight. Following AMP1, AMP2 and AMP3 treatment, the sections were incubated with Opal 570 (AKOYA BIOSCIENCES, Cat# FP1488001KT) for C1, Opal 620 (AKOYA BIOSCIENCES, Cat# FP1495001KT) for C2 and Opal 520 (AKOYA BIOSCIENCES, Cat# FP1487001KT) for C3 in 1:1000 dilution at RT, for 15’. Then slides were mounted with VECTASHIELD® HardSet™ Antifade Mounting Medium with DAPI (Maravai LifeSciences, Cat# H1500). Images were then captured using a Zeiss Axio Observer microscope with an ApoTome (Carl Zeiss Corp.).

### scRNA-seq data analysis

We utilized Cell Ranger (v 3.1.0, from 10x Genomics) to align reads of scRNA-seq data to the mouse reference genome (mm10). All the samples passed quality control measures for Cell Ranger (Table 1), and the filtered gene-barcode matrices were used for the following analyses. For *Mesp1^Cre^* four time point dataset analyses, Scran v1.10.2 was used to normalize the individual data sets by “computeSumFactors” function with the deconvolution method for scaling normalization^71^. The cells were clustered by densityClust v0.3 with the density peak clustering algorithm ^21^. We found batch effects exist in the scRNA-seq data sets from different time points (*Mesp1^Cre^* data) or experimental perturbations (*Mesp1^Cre^ Tbx1* or *Tbx1^Cre^ Tbx1* Ctrl vs cKO data). Therefore, we performed batch corrections before we comprehensively analyzed gene expression values across these scRNA-seq data sets. We employed the MNN (mutual nearest neighbors) method to identify shared cell types across data sets, and corrected batches, according to the shared cell types, by the MNN batch correction method^20^. More specifically, we removed batch effects by “fastMNN” function of the Scran package, with input of the normalized counts from individual datasets. The *Mesp1^Cre^* Ctrl data from different stages are comprised of homogeneous cell types (transcriptomes of each cell type are concordant across datasets), but the *Mesp1^Cre^ Tbx1* Ctrl with cKO and then separately, the *Tbx1^Cre^* Ctrl versus cKO, are comprised of heterogeneous cell types (the same cell types from different conditions, Ctrl vs cKO, with dissimilar gene expression). Thus, we aligned gene expression values for Ctrl vs cKO data using the RPCI (reference principal component integration) method which utilizes the global gene reference to calibrate the gene expression changes of heterogeneous cell types^72^. In detail, we combined individual data sets by the “scMultiIntegrate” function of the RISC package and outputted the corrected gene expression values after the clustering by Seurat 3.1.5^73^. Then, the corrected data sets were processed on Scanpy v1.4.3^74^ for cell trajectory analysis, with the PAGA approach^28^. The RISC software^72^ was also used to identify differentially expressed genes of *Tbx1* Ctrl vs cKO embryos in each cluster by a Negative Binomial generalized linear model^75^ (“scDEG” function of RISC package), with adjustive p-values < 0.05 and logFC (log fold change) > 0.25 or <−0.25. A customized R script was used for comparison of gene lists. The clusterProfiler v3.10.1^76^ was used for Gene Ontology pathway analysis, with adjustive P-values < 0.05.

The scRNA-seq data can be viewed at this URL: https://scviewer.shinyapps.io/heartMLP

### ATAC-seq data analysis

ATAC-seq analysis pipeline has been described previously^66^.

We removed Nextera Transposase Sequence primers^77^ in the range 33 to 47 bp using cutadapt^78^ with the following option -a CTGTCTCTTATACACATCTCCGAGCCCACGAGAC -A CTGTCTCTTATACACATCTGACGCTGCCGACGA. Sequences were then aligned to the mouse genome (mm9) using Bowtie2 2.3.4.3 (Langmead and Salzberg, 2012) with default parameters. Only uniquely mappable reads were retained. A customized R script was used to remove reads with mates mapping to different chromosomes, or with discordant pairs orientation, or with a mate-pair distance >2 kb, or PCR duplicates (defined as when both mates are aligned to the same genomic coordinates). Reads mapping to the mitochondrial genome were also removed. ATAC peaks in each sample were identified using MACS2 2.1.2.1^79^ with the option -- nomodel --shif100 --extsize 200. The differentially enriched regions (DARs) of *Tbx1^Cre^* Ctrl vs cKO were obtained using DiffBind 2.14.0^80^ by loading all of the MACS2 peaks, default parameters were used except for dba.count method were summits has been set to 250. After that, we formed a consensus list of enriched regions for each condition using the intersectBed function from the BedTools 2.29^81^ with the default minimum overlap and retaining only the peak regions common to all three replicates. We defined common peaks the regions that were common to the consensus peaks of both *Tbx1^Cre^* Ctrl and cKO (using again the intersectBed function). The common peaks in*Tbx1^Cre^* Ctrl vs cKO data that did not intersect the DARs were defined as common accessible regions (CARs). Both DARs and CARs were filtered out by removing peaks intersecting blacklist regions (Encode mm9 blackregions Version 2) using findOverlapsOfPeaks of ChIPpeakAnno v3.22.4^82^ before any further analysis. Then DARs or CARs were filtered with the consensus *Mesp1^Cre^* ATAC-seq peaks using intersectBed.

Transcription factor binding motifs were obtained using the findMotifsGenome program of the HOMER suite^83^. For peak annotation as *cis-*regulatory regions, GREAT^42^ was used with default settings. The comparison of the gene list from DARs and DEGs was performed using standard R-scripts. The clusterProfiler v3.10.1 was used for Gene Ontology pathway analysis. Coverage heat-maps and average enrichment profiles (TSS +/− 10Kb) in each experimental condition were obtained using ngs.plot^84^ or deepTools2^85^.

### ChIP-seq data analysis

We removed adapters using cutadapt^78^ with -a option and a set of adapters detected with FASTQC ^86^. Sequences were then trimmed using TrimGalore with option -length 0. Then sequences were aligned to the mouse genome (mm9) using Bowtie2 2.3.4. with default parameters. Only uniquely mappable reads were retained. A customized R script was used to remove reads with mates mapping to different chromosomes, or with discordant pairs orientation, or with a mate-pair distance >2 kb, or PCR duplicates (defined as when both mates are aligned to the same genomic coordinate). ChIP-seq peaks in each sample were identified using MACS2 2.1.2.1 with default parameters. Then, a consensus list of enriched regions was obtained using the intersectBed function from the BedTools 2.29 with the default minimum overlap and retaining only the peak regions common to at least two out of the three replicates. Peaks were filtered by removing those overlapping with blacklist regions (Encode mm9 black regions Version 2) using findOverlapsOfPeaks of ChIPpeakAnno. The transcription factor binding motifs were obtained using the findMotifsGenome program with -size given parameter of the HOMER suite. For peak annotation as *cis-*regulatory regions, GREAT was used with the default settings. The comparison of the gene lists of DAR, DEG and ChIP regions was performed using standard R-scripts. IGV 2.4.8^85^ was used for peak visualization. Coverage heat-maps and average enrichment profiles (TSS +/− 10Kb) in each experimental condition were obtained using ngs.plot or deepTools2. Significance of the overlap between two list of peaks was evaluated using the ChIPseeker enrichPeakOverlap using mm9 annotation.

## Data and Code Availability

The datasets generated during this study are available at GEO repository under the accession number GSE158941. The scRNA-seq data can be viewed at this URL: https://scviewer.shinyapps.io/heartMLP

## Quantification and Statistical Analysis

Specific statistical tests were described in the Results and Figure legends.

## Acknowledgements

We thank Genomics core, especially David Reynolds, Director of the Genomics Core, Shahina Maqbool, Director of the Epigenomics core as well as the Flow Cytometry and Gene modification facilities at Einstein. We appreciate all the help in data analysis and training by Masako Suzuki at Einstein. We thank Drs. Bin Zhou and Peter Scambler for reading this manuscript and providing helpful suggestions. We also thank NYU Center for Genomics and Systems Biology Genomics Core and NYU Langone’s Genome Technology Center is partially supported by the Cancer Center Support Grant P30CA016087 at the Laura and Isaac Perlmutter Cancer Center, especially Peter Meyn. This work was supported by grants from the National Institutes of Health P01HD070454 (BEM, DZ), R01HL153920 (BEM, DZ), R01HD096770 (LC) and R01HL108643 (LC). The work was also supported by a grant from the Foundation Leducq (Transatlantic Network of Excellence 15CVD01; BEM, LC, RGK, AB) and the Agence Nationale pour la Recherche Heartbox project (RGK). Dr. Nomaru was supported by an American Association Grant (19POST34380281).

## Author Contributions

Conceptualization: H.N. and B.E.M

Validation: H.N., B.E.M

Formal Analysis, H.N., Y.L., C.D., D.R., A.C., W.W., C.A and D.Z

Investigation, H.N., W.W., S.R. and A.D.;

Data curation, H.N., Y.L., D.R., A.C. Writing-Original Draft, H.N. and B.E.M.

Writing-Review & Editing, H.N, Y.L., C.D., D.R., A.C., C.A., L.C., R.G.K., A.B., D.Z. and B.E.M.

Visualization, H.N., Y.L., C.D., D.R., A.C. Supervision, D.Z. and B.E.M

Funding Acquisition, H.N., L.C., R.G.K., A.B. and B.E.M. Resources, C.C

---

**Extended Data Fig 1. Sample preparation and cluster identification of *Mesp1*+ lineages at E8.0 to E10.5 for the scRNA-seq dataset**

**A.** Scheme of sample preparation for scRNA-seq. After dissection of tissues from the embryo, GFP+ cells were sorted. Following cell counting with checking cell viability, cells were loaded to 10x Chromium system.

**B.** The gating example of FACS purification. Only the GFP+, DAPI-population was collected for scRNA-seq.

**C.** Heatmap of average gene expression level of marker genes in all clusters of *Mesp1*+ lineages at E8.0 to E10.5 in the scRNA-seq dataset.

**D.** tSNE plot colored by clusters with the cluster selection information. Red dotted line represents the cluster group used for further analysis in Figure 2A.

**E.** tSNE plot colored by expression level of marker genes. The color spectrum from grey to red indicates expression level from low to high.

**Extended Data Fig 2. Gene expression on single-cell embedding graph of *Mesp1*+ lineages at E8.0 to E10.5 in the scRNA-seq dataset**

**A.** Single-cell embedding graph separated by stages (E8.0, E8.25, E9.5 and E10.5), colored by clusters. Cluster colors are consistent with those in Figure 2A.

**B.** Single-cell embedding graph separated by stages, colored by pseudotime. The color spectrum from red/orange is the early pseudotime point to blue/purple is late pseudotime point.

**C.** Single-cell embedding graph of the CPM lineages separated by stages, colored by clusters. Cluster colors are consistent with those in Figure 2D.

**D.** Single-cell embedding graph colored by expression level. The color spectrum from blue, then green to yellow indicates expression levels from low to high. Grey indicates no expression. The genes are consistent with Figure 2F.

**E.** Heatmap of average gene expression of the genes enriched in the cluster selected for the single-cell embedding graph. The genes are consistent with Figure 2G.

**F.** Single-cell embedding graph colored by expression level of the marker genes of BrM, aSHF and pSHF in all clusters (top), CPM (middle) and MLPs (bottom). The color spectrum from blue, then green to yellow indicates expression levels from low to high. Grey indicates no expression.

**G.** Serial RNAscope *in situ* hybridization pictures with *Nrg1*, *Aplnr* and *Isl1* mRNA probes on sagittal sections E8.5 (n = 3). All the sections are from same embryo as Figure 2H. Green, *Nrg1*; Red, *Aplnr*; Blue, *Isl1*.

**H.** Serial RNAscope *in situ* hybridization with *EGFP*, marking *Mesp1*+ lineage cells, *Tbx1* and *Isl1* mRNA probes on sagittal sections from *Mesp1^Cre^;ROSA26-GFP*^*flox*/+^ embryos at E8.5 (n = 3). All the sections are from same embryo as Figure 2K. Green, *EGFP*; Red, *Tbx1*, Blue, *Isl1*.

**I.** Serial RNAscope *in situ* hybridization pictures with *Nrg1*, *Aplnr* and *Isl1* mRNA probes on sagittal sections E9.5 (n = 3). All the sections are from same embryo as Figure 2H. Green, *Nrg1*; Red, *Aplnr*; Blue, *Isl1*.

**J.** Serial RNAscope *in situ* hybridization with *EGFP*, marking *Mesp1*+ lineage cells, *Tbx1* and *Isl1* mRNA probes on sagittal sections from *Mesp1^Cre^;ROSA26-GFP*^*flox*/+^ embryos at E9.5 (n = 3). All the sections are from same embryo as Figure 2K. Green, *EGFP*; Red, *Tbx1*, Blue, *Isl1*. Abbreviations: heart (ht), outflow tract (oft), pharyngeal arch (PA), pharynx (ph), right ventricle (RA), venous pole (vp), 1, 2 and 3 indicate the first, second and third pharyngeal arches, respectively.

**Extended Data Fig 3. Cluster identification of *Mesp1^Cre^;Tbx1* Ctrl and cKO at E9.5 datasets, *Tbx1^Cre^;Tbx1* Ctrl and cKO at E8.5 and E9.5 datasets**

**A.** Single-cell embedding graph with PAGA plot of the integrated dataset from *Mesp1*^*Cre*/+^;*ROSA26-GFP*^*flox*/+^ (Ctrl) vs *Mesp1*^*Cre*/+^;*ROSA26-GFP*^*flox*/+^;*Tbx1*^*flox/flox*^ (cKO) embryos at E9.5, colored by cluster. C0: Sk/L, skeleton/limb; C1: BrM, branchiomeric muscle progenitors; C2: Lung PC, lung progenitors; C3: ST, septum transversum; C4: MLP, multilineage progenitors; C5: aSHF/SoM, anterior CPM and somatic mesoderm; C6: MCs, mesenchyme cells; C7: pSHF, posterior CPM; C8: CMs, cardiomyocyte progenitor cells; C9: PEO, proepicardial organ; C10: Sk/L, skeleton/limb; C11: pSHF, posterior CPM; C12: Lung PC, lung progenitor cells; C13: Endo, endocardium and endothelial cells; C14: Endo, endocardium and endothelial cells; C15: Blood, blood cells; C16: Endo, endocardium and endothelial cells; C17: ST, septum transversum; C18: CMs, cardiomyocyte progenitor cells; C19: Endothelial, endothelial cells.

**B.** Single-cell embedding graph separated by genotype, colored by clusters. Cluster colors are consistent with those in Extended Data Fig3A.

**C.** Single-cell embedding graph of CPM populations colored by pseudotime. The color spectrum from red/orange is the early pseudotime point to blue/purple is late pseudotime point.

**D.** Single-cell embedding graph colored by expression level, related to Figure 3F. The color spectrum from blue to yellow indicates expression levels from low to high. Grey indicates no expression.

**E.** Single-cell embedding graph colored by expression level of the marker genes. The color spectrum from blue to yellow indicates expression levels from low to high. Grey indicates no expression.

**F.** Single-cell embedding graph with PAGA plot of the integrated dataset from *Tbx1*^*Cre*/+^;*ROSA26-GFP*^*flox*/+^ (Ctrl) vs *Tbx1*^*Cre/flox*^; *ROSA26-GFP*^*flox*/+^ (cKO) embryos at E8.5 and E9.5, colored by cluster. C0: MCs, mesenchyme; C1: MCs, mesenchyme; C2: MCs, mesenchyme; C3: BrM, branchiomeric muscle progenitor cells; C4: aSHF/SoM, anterior CPM and somatic mesoderm; C5: MLP, multi-lineage progenitor cells; C6: Endo, endocardium and endothelial cells; C7: MCs, mesenchyme; C8: Blood, blood cells; C9: MCs, mesenchyme; C10: Epi, epithelia including endoderm and ectoderm; C11: Epi, epithelia including endoderm and ectoderm; C12: pSHF, posterior CPM; C13: CMs, cardiomyocyte progenitor cells; C14: OV, otic vesicle; C15: NPG, cochlear-vestibular ganglion neural progenitor cells; C15: NPG, cochlear-vestibular ganglion neural progenitor cells; C17: Epi, epithelia including endoderm and ectoderm; C18: OV, otic vesicle.

**G.** Single-cell embedding graph separated by genotype and stage, colored by clusters. Cluster colors are consistent with those in Extended Data Fig3F.

**H.**Single-cell embedding graph of CPM clusters colored by pseudotime. The color spectrum from red/orange is early pseudotime point to blue/purple as late pseudotime point.

**I.** Single-cell embedding graph colored by expression level, related to Figure 3J. The color spectrum from blue to yellow indicates expression level from low to high. Grey indicates no expression.

**J.** Single-cell embedding graph colored by expression level of marker genes. The color spectrum from blue to yellow indicates expression levels from low to high. Grey indicates no expression.

**Extended Data Fig 4. Differentially expressed genes analysis in CPM lineages**

**A-D**. Comparison of differentially expressed genes (DEGs) from *Mesp1*^*Cre*/+^;*ROSA26-GFP*^*flox*/+^ (Ctrl) vs *Mesp1*^*Cre*/+^;*ROSA26-GFP*^*flox*/+^;*Tbx1*^*flox/flox*^ (cKO) embryos at E9.5 and *Tbx1*^*Cre*/+^;*ROSA26-GFP*^*flox*/+^ (Ctrl) vs *Tbx1*^*Cre/flox*^;*ROSA26-GFP*^*flox*/+^ (cKO) in BrM (A), aSHF/SoM(B), pSHF-7, cluster 7 (C), and CMs (D). X-axis indicates log2-fold change of *Mesp1^Cre^* DEGs. Y-axis indicates log2-fold change of *Tbx1^Cre^* DEGs. Each dot indicates a gene (*adj. p-value* < *0.05*), and red indicates significant fold change (|fold change|> 0.25) in both comparisons. As in Extended Data Fig3A, there are two pSHF clusters (C7 and C11) in the *Mesp1^Cre^* dataset. So, we compared *Mesp1^Cre^* cluster 7 DEG vs *Tbx1^Cre^* cluster 12 DEG (C) and *Mesp1^Cre^* cluster 11 DEG vs *Tbx1^Cre^* cluster 12 DEG, respectively. Then we merged two DEGs as pSHF DEGs.

**E-I**. Enriched biological processes based upon GO and related genes found in downregulated (E) and upregulated (F) genes in BrM, downregulated genes in aSHF/SoM clusters (G), upregulated genes in pSHF (H) and upregulated genes in CMs (I) of *Tbx1* cKO data. X-axis indicates adjusted p-values. There was no enriched GO found in upregulated genes in aSHF/SoM, downregulated gene in pSHF and CMs of *Tbx1* cKO data.

**J.** Single-cell embedding graph of the CPM lineages from *Mesp1^Cre^;Tbx1* Ctrl and cKO at E9.5 datasets and *Tbx1^Cre^;Tbx1* Ctrl and cKO at E9.5 separated by genotype and colored by expression level. The genes are found as downregulated in MLP and other CPM clusters (Figure 4B). The color spectrum from blue to yellow indicates expression levels from low to high. Grey indicates no expression.

**K.** Violin plot of the genes in each CPM cluster from *Mesp1^Cre^;Tbx1* Ctrl and cKO data at E9.5. The genes are found as downregulated in MLPs and other CPM clusters (Figure 4B).

**L.** Violin plot of the genes in each CPM cluster from *Tbx1^Cre^;Tbx1* Ctrl and cKO data at E8.5 and E9.5. The genes found that were downregulated in MLPs and other CPM clusters in data from mutant embryos (Figure 4B).

**Extended Data Fig 5. Gene expression of DEGs in CPM lineages, related to Figure 4**

**A.** Violin plot of *Aplnr*, *Nrg1*, *Pax8* and *Bdnf* expression in MLPs from *Mesp1^Cre^;Tbx1* Ctrl and cKO data at E9.5.

**B.** Violin plot of *Aplnr*, *Nrg1*, *Pax8* and *Bdnf* expression in MLP from *Tbx^Cre^;Tbx1* Ctrl and cKO at data at E8.5 and E9.5.

**C.** Heatmap of expression of *Aplnr*, *Nrg1*, *Pax8*, *Bdnf* and *Pax6* in MLPs from *Mesp1^Cre^;Tbx1* Ctrl and cKO data at E9.5. Row indicates the expression of each cell.

**D.** Heatmap of expression of *Aplnr*, *Nrg1*, *Pax8*, *Bdnf* and *Pax6* in MLPs from *Tbx^Cre^;Tbx1* Ctrl and cKO data at E8.5 and E9.5. Row indicates the expression of each cell.

**E.** RNAscope *in situ* hybridization with *EGFP*, *Aplnr* and *Pax8* mRNA probes on transverse section from *Mesp1*^*Cre*/+^;*ROSA26-GFP*^*flox*/+^;*Tbx1*^*flox*/+^ (Het, Ctrl) embryos (upper panel; n = 3) and *Mesp1*^*Cre*/+^;*ROSA26-GFP*^*flox*/+^;*Tbx1*^*flox/flox*^ (cKO) embryos (lower panel; n = 3) at E9.5. Green; *GFP*, Red; *Aplnr*, Blue; *Pax8*. White dotted line indicates the position in higher magnification images.

**F.** Higher magnification images of Extended Data Fig5E. Upper left, merged image; Upper right, green channel (*GFP*), Lower left, red channel (*Aplnr*); Lower right, blue channel (*Pax8*). Scale bar; 100 μm. Dotted line indicates the boundary of ectoderm and mesoderm.

**G.** RNAscope *in situ* hybridization with *GFP*, *Aplnr* and *Pax8* mRNA probes on transverse section from *Tbx1*^*Cre*/+^;*ROSA26-GFP*^*flox*/+^ (Het, Ctrl) embryos (upper panel; n = 3) and *Tbx1*^*Cre/flox*^;*ROSA26-GFP*^*flox*/+^ (cKO) embryos (lower panel; n = 3) at E9.5. Green; *GFP*, Red; *Aplnr*, Blue; *Pax8*. White dotted line indicates the position in higher magnification images.

**H.** Higher magnification images of Extended Data Fig5E. Upper left, merged image; Upper right, green channel (*EGFP*), Lower left, red channel (*Aplnr*); Lower right, blue channel (*Pax8*) Scale bar; 100 μm. Dotted line indicates the boundary of ectoderm and mesoderm. Abbreviations: atrium (a), aorta (ao), ectoderm(e), neural tube (nt), pharynx (ph), ventricle (v).

**Extended Data Fig 6. ATAC-seq data analysis, related to Figure 5**

**A.** Average ATAC-seq read densities at regions ±3kb from transcription start sites of GFP+ cells from *Mesp1*^*Cre*/+^; *ROSA26-GFP*^*flox*/+^ (Left), *Tbx1*^*Cre*/+^; *ROSA26-GFP*^*flox*/+^ (Middle) and *Tbx1*^*Cre/flox*^; *ROSA26-GFP*^*flox*/+^ (Right).

**B.** Heatmap of ATAC-seq read densities at regions ±3kb from transcription start sites of GFP+ cells from *Mesp1*^*Cre*/+^; *ROSA26-GFP*^*flox*/+^ (Left), *Tbx1*^*Cre*/+^; *ROSA26-GFP*^*flox*/+^ (Middle) and *Tbx1*^*Cre/flox*^; *ROSA26-GFP*^*flox*/+^ (Right).

**C.** Pie chart of genome distribution of ATAC-seq peaks of GFP+ cells from *Mesp1*^*Cre*/+^; *ROSA26-GFP*^*flox*/+^ (Left), *Tbx1*^*Cre*/+^; *ROSA26-GFP*^*flox*/+^ (Middle) and *Tbx1*^*Cre/flox*^; *ROSA26-GFP*^*flox*/+^ (Right).

**D.** Volcano plot of the ATAC-seq peaks from *Tbx1^Cre^ Tbx1* Ctrl vs *Tbx1^Cre^ Tbx1* cKO dataset. The pink dots indicate differentially accessible regions (FDR ≤ 0.05)

**E.** Pie chart of genome distribution of CAR-Mesp1.

**F.** Pie chart of genome distribution of DAR-Mesp1.

**Extended Data Fig 7. ChIP-seq data analysis**

**A.** Scheme of TBX1 *in vivo* ChIP-seq with Avi-tag. The Avi-tag sequence was inserted in 3’ end of *Tbx1* exon 7 (Left). In *Tbx1-Avi;BirA* homozygous mouse, biotin was bind to Avi-tag of the TBX1 protein (Right).

**B.** Average ChIP read densities of each replicate (Tbx1-ChIP; top, Input; bottom) at regions from transcription start sites (TSS) to transcription end site (TES).

**C.** Venn diagram of peaks from three replicates of ChIP-seq. We used the peaks found in at least two replicates.

**D.** The known motifs found in ChIP-seq regions by Homer software.

**E.** The novel motifs found in ChIP-seq regions by Homer software.

**F.** Chromatin accessibility of ChIP-seq peak regions in three replicates of ATAC-seq data. Left, ATAC-seq data from *Tbx1^Cre^ Tbx1* Ctrl, Right, ATAC-seq data from *Tbx1^Cre^ Tbx1* cKO data.

**Supplementary Table 1. Marker genes of cell clusters *in Mesp1^Cre^* four timepoint dataset, related to Figure 1.**

**Supplementary Table 2. Marker genes of cell clusters *in Mesp1^Cre^ Tbx1* Ctrl and cKO at E9.5 dataset, related to Figure 3.**

**Supplementary Table 3. Marker genes of cell clusters *in Tbx1^Cre^ Tbx1* Ctrl and cKO at E8.5 and E9.5 dataset, related to Figure 3.**

**Supplementary Table 4. Differentially expressed genes (adjusted p-value < 0.05) in each cluster of *Mesp1^Cre^ Tbx1* Ctrl and cKO at E9.5 dataset, related to Figure 4.**

**Supplementary Table 5. Differentially expressed genes (adjusted p-value < 0.05) in each cluster of *Tbx1^Cre^ Tbx1* Ctrl and cKO at E9.5 dataset, related to Figure 4.**

**Supplementary Table 6. Differentially expressed genes (adjusted p-value < 0.05, fold change > |0.25| in both *Mesp1^Cre^ Tbx1* Ctrl vs cKO at E9.5 data and *Tbx1^Cre^ Tbx1* Ctrl and cKO at E9.5 data) in MLP of *Tbx1* Ctrl vs cKO at E9.5, related to Figure 4.**

**Supplementary Table 7. Differentially expressed genes (adjusted p-value < 0.05, fold change > |0.25| in both *Meps1^Cre^ Tbx1* Ctrl vs cKO at E9.5 data and *Tbx1^Cre^ Tbx1* Ctrl and cKO at E9.5 data) in BrM, aSHF/SoM, pSHF and CMs of *Tbx1* Ctrl vs cKO at E9.5, related to Figure 4.**

**Supplementary Table 8. Gene ontology biological pathway found in DEGs of MLP, related to Figure 4.**

**Supplementary Table 9. Gene ontology biological pathway found in downregulated in DEGs of CPM cells (BrM and aSHF/SoM), related to Figure 4.**

**Supplementary Table 10. Gene ontology biological pathway found in upregulated in DEGs of CPM cells (BrM pSHF, CMs), related to Figure 4.**

**Supplementary Table 11. DARs-Mesp1 ATAC-seq peak regions with annotated genes, related to Figure 5.**

**Supplementary Table 12. The motif enriched in DARs-Mesp1 ATAC-seq peak regions, related to Figure 5.**

**Supplementary Table 13. TBX1 ChIP-seq peak regions with annotated genes, related to Figure 6.**

**Supplementary Table 14. The motif enriched in TBX1 ChIP-seq peak regions, related to Figure 6.**

