## Supplemental Figures for "Single cell multi-omic analysis identifies a *Tbx1*-dependent multilineage primed population in the murine cardiopharyngeal mesoderm"

#### Supplement figure 1

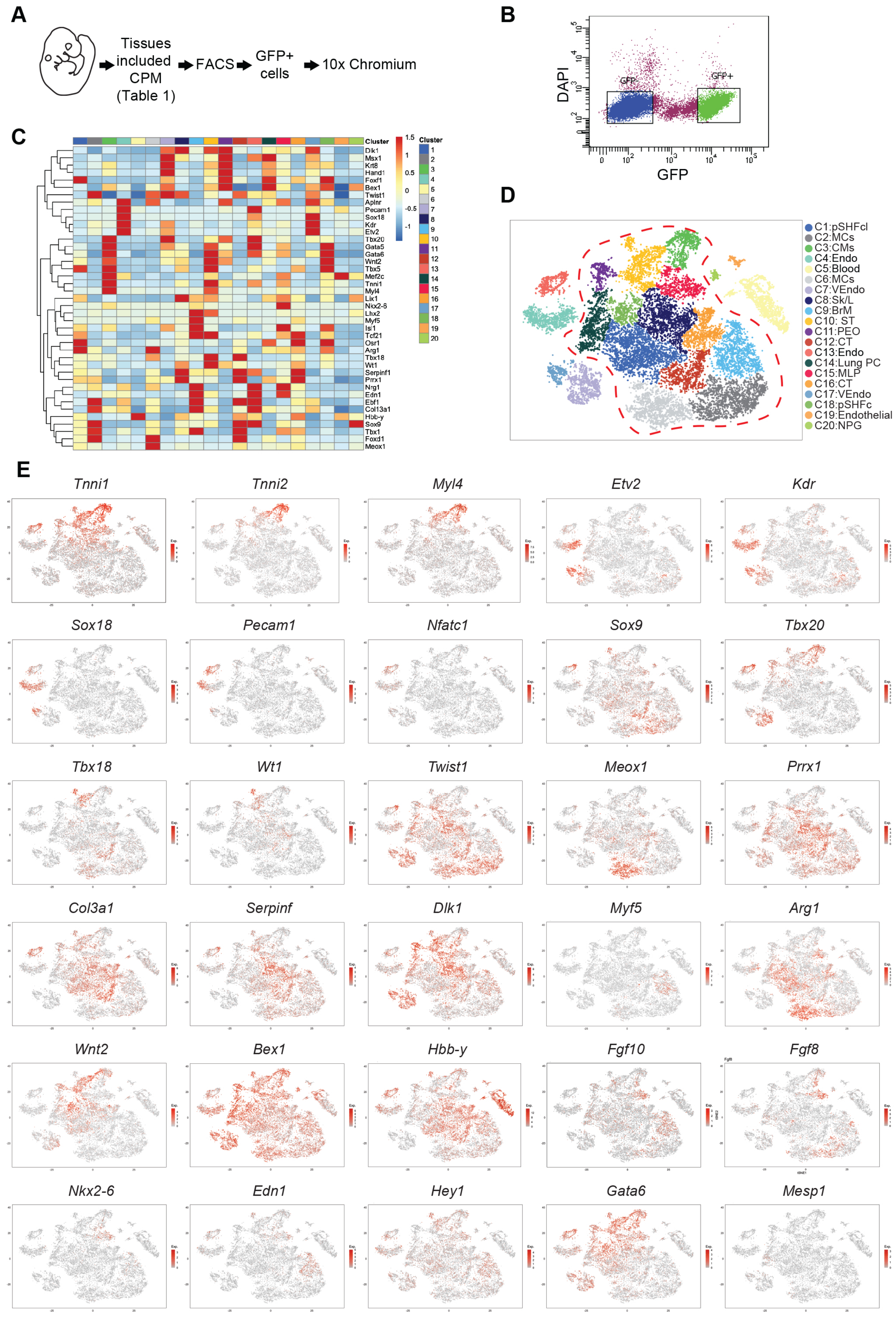

### Supplement figure 2

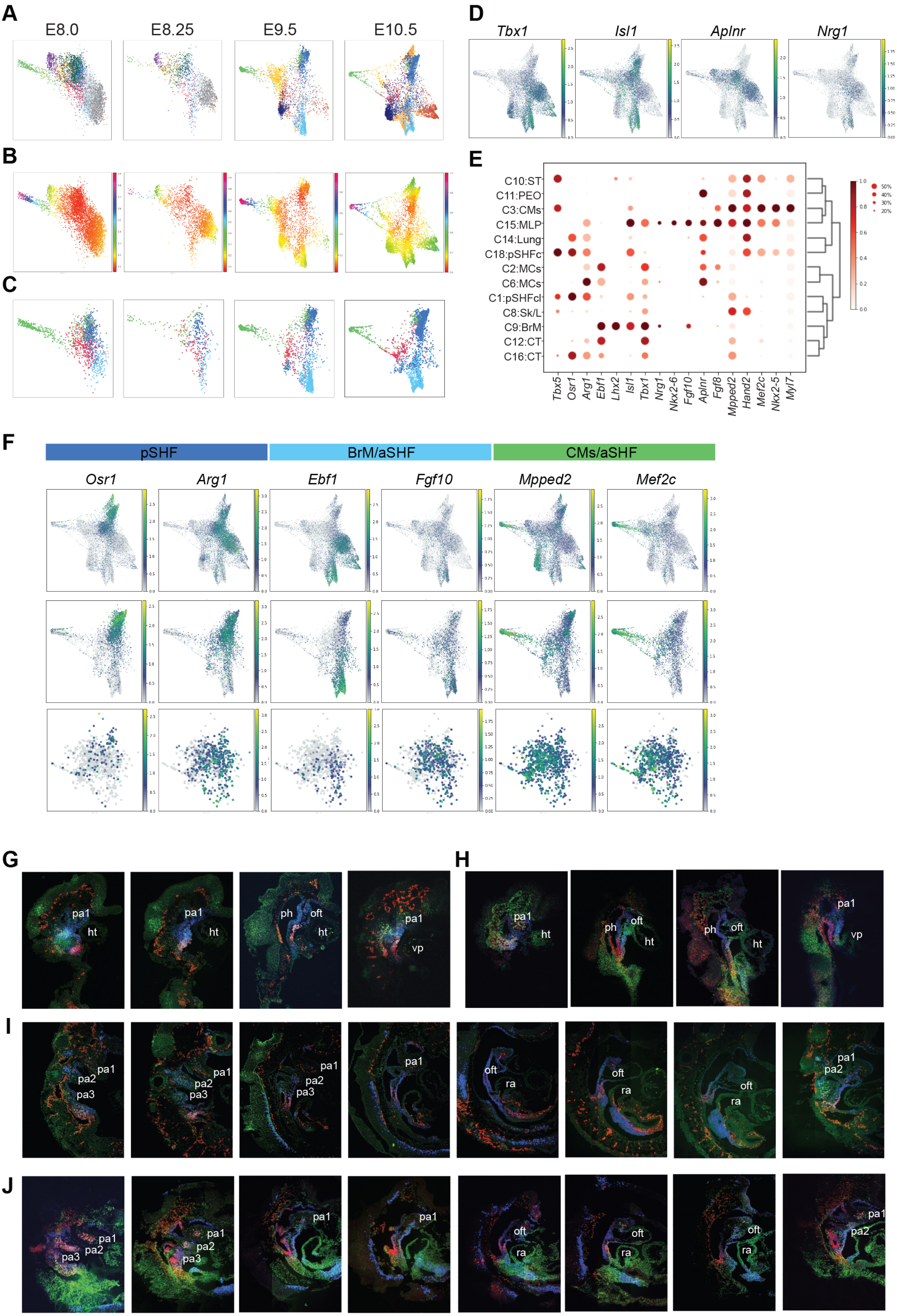

### Supplement figure 3

#### *Mesp1^Cre* lineage

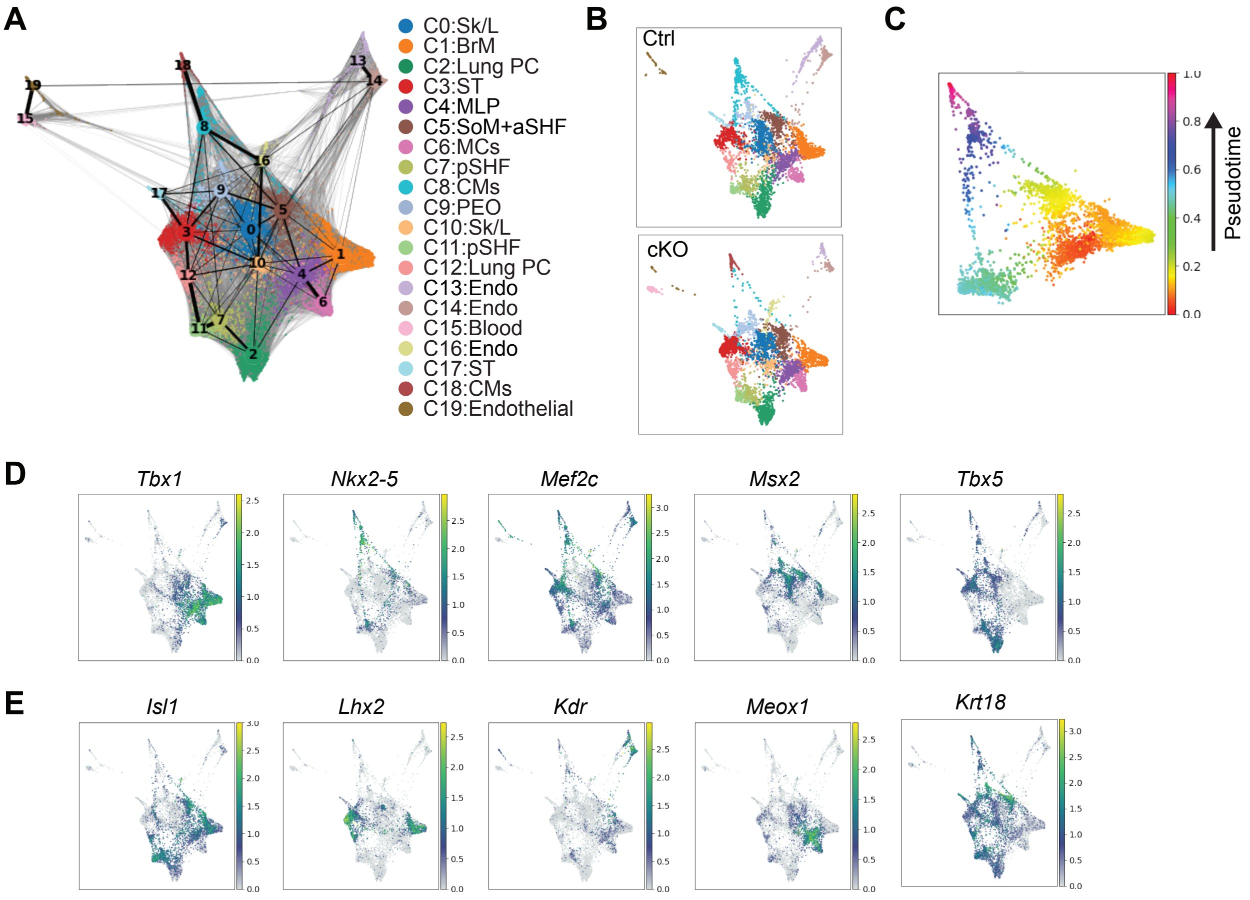


#### *Tbx1^Cre* lineage

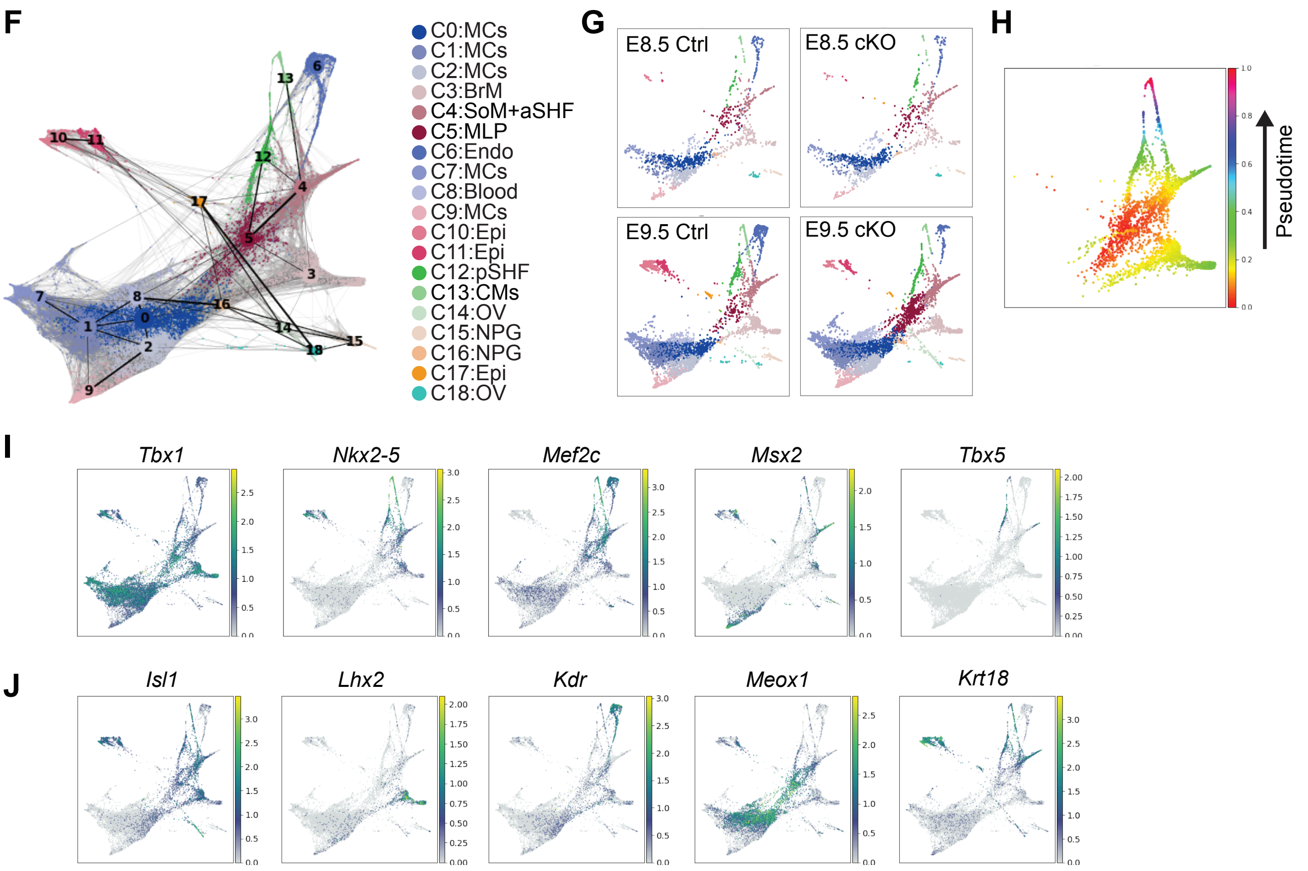

### Supplement figure 4

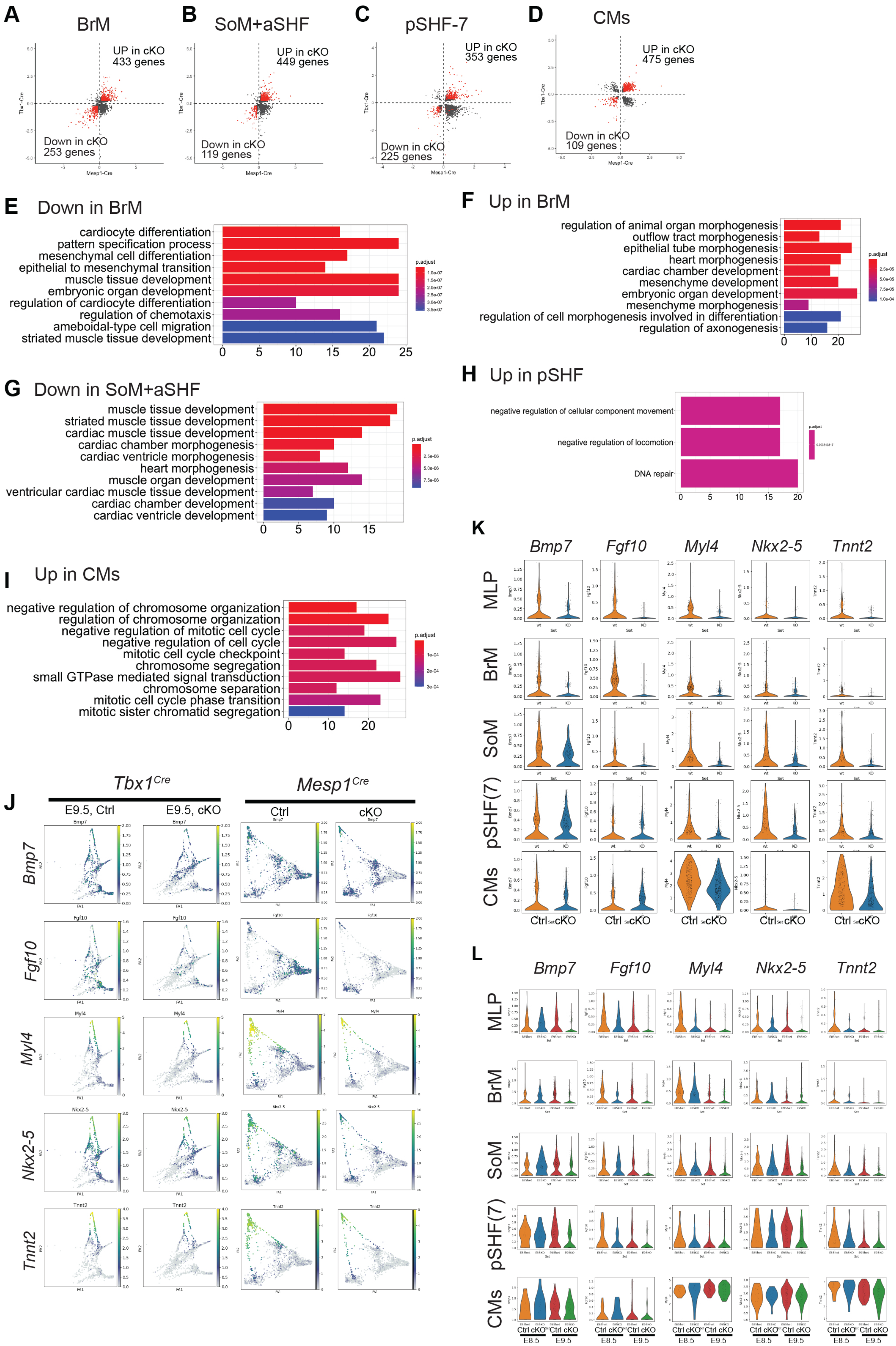

### Supplement figure 5

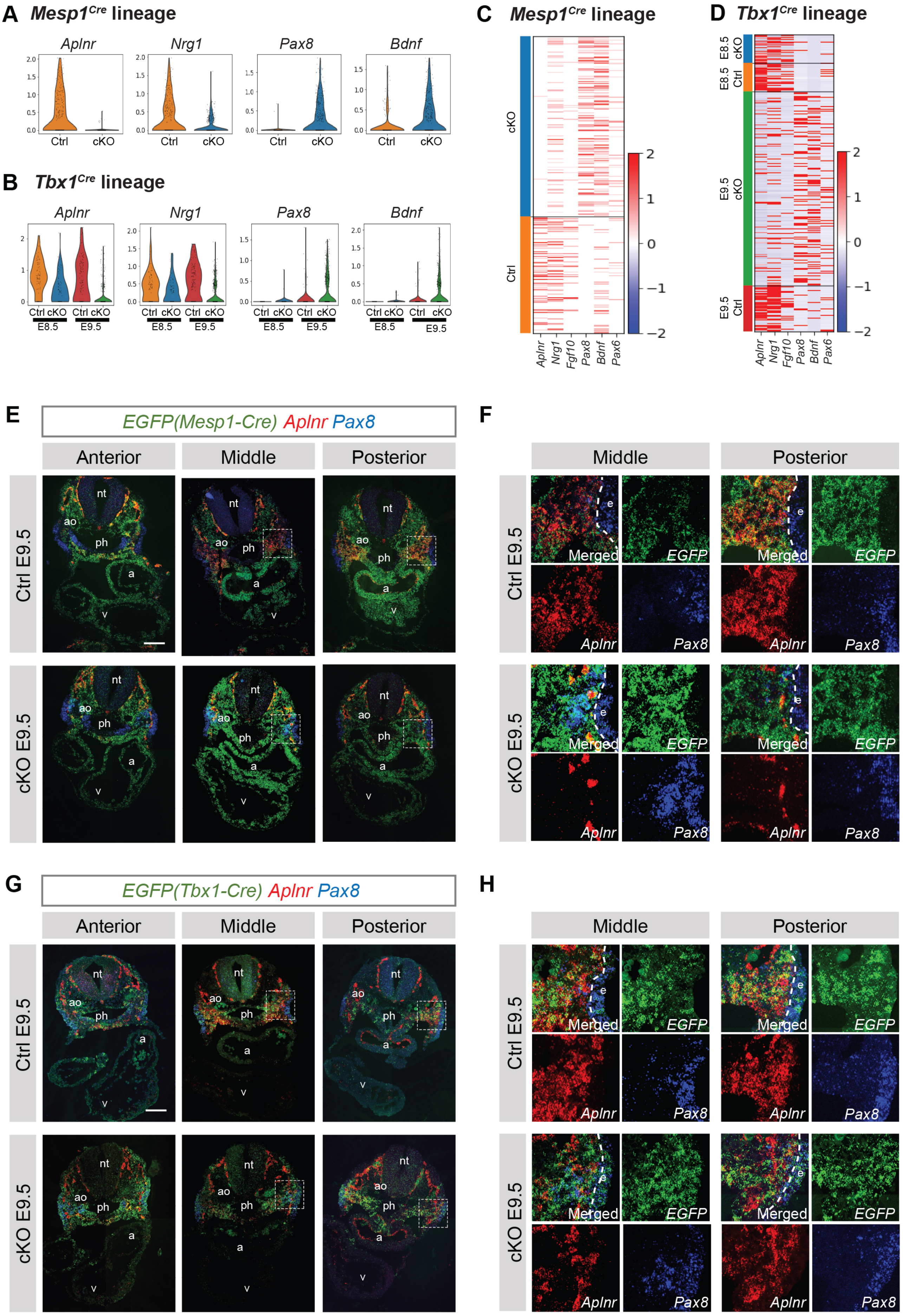

#### Supplement figure 6

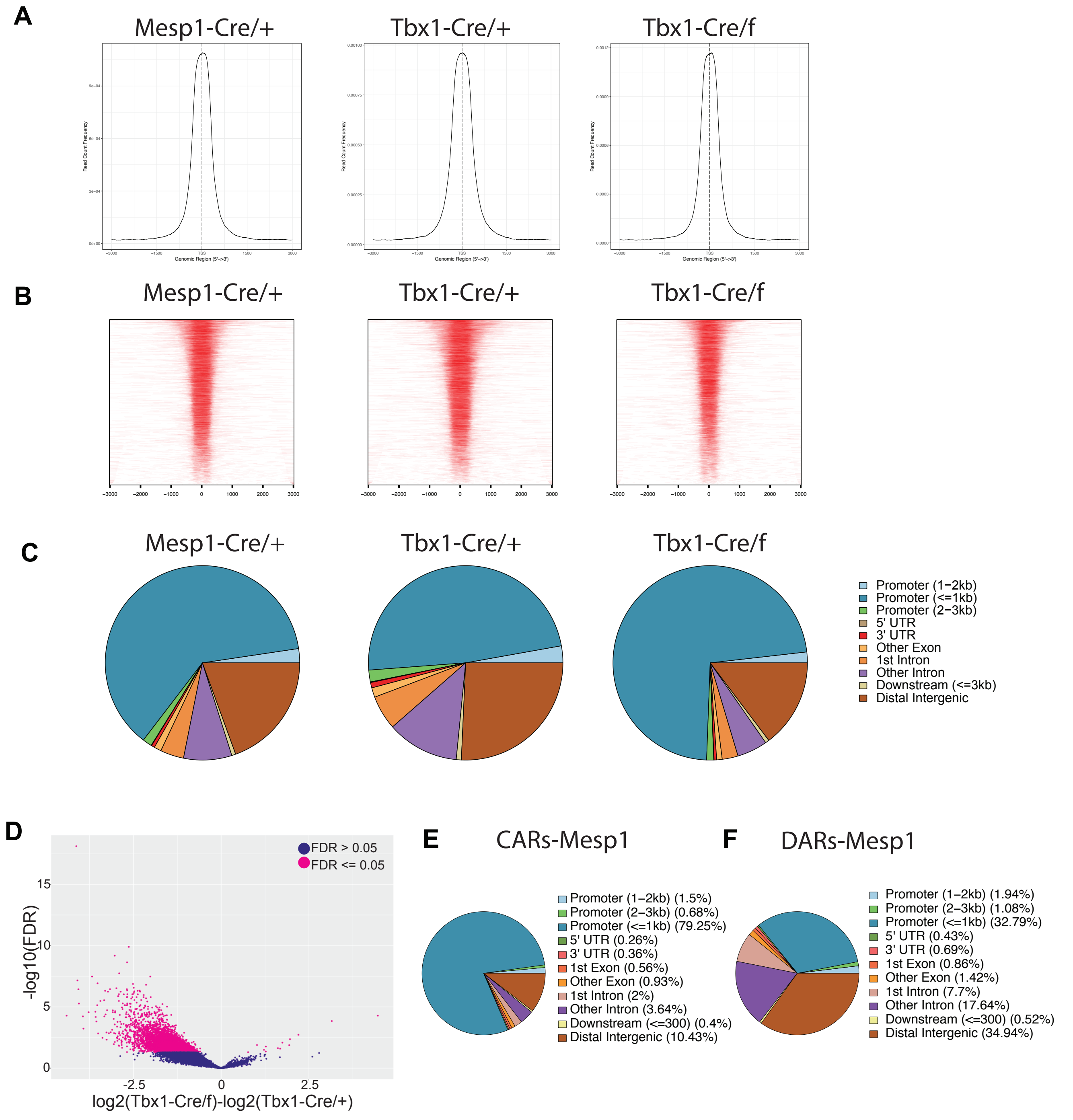

#### Supplement figure 7

**A** *Tbx1 3'-Avi*

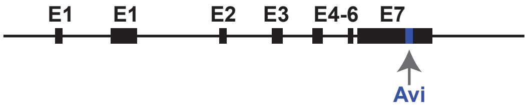


*Tbx1-Avi;BirA* homozygous mouse

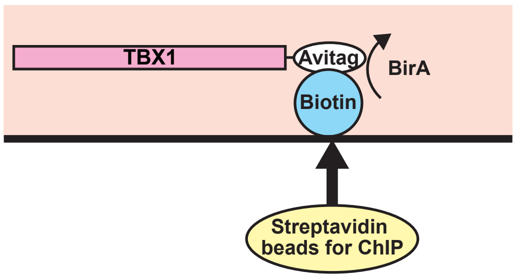


**B**

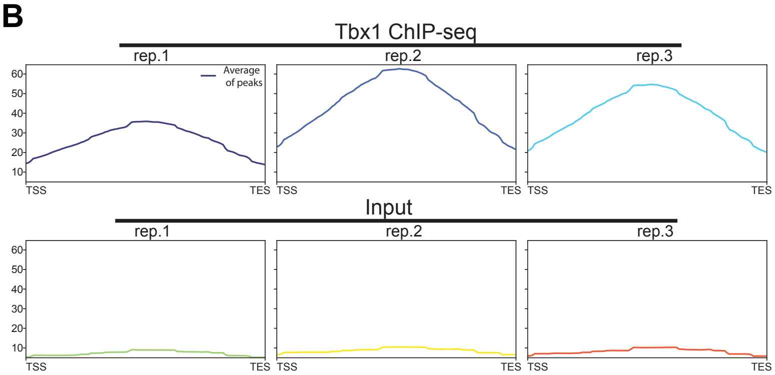


**C**

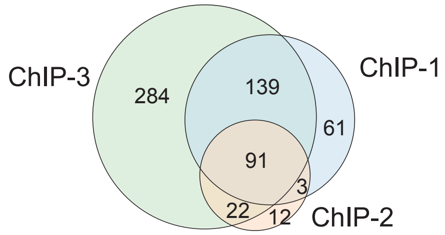


**D** known

| Rank | Motif | Transcription factor | p-value | Back ground | Targets |
|---|---|---|---|---|---|
| 1 | 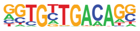 | Tbx20(T-box) | 1e-59 | 4.44% | 37.25% |
| 2 | 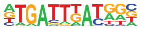 | PBX2(Homeobox) | 1e-36 | 13.08% | 45.88% |
| 3 | 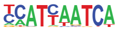 | Pdx1(Homeobox) | 1e-34 | 16.16% | 49.80% |
| 4 | 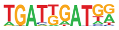 | HOXA1(Homeobox) | 1e-33 | 5.47% | 29.41% |

**E** *De novo*

| Rank | Motif | Transcription factor | p-value | Back ground | Targets |
|---|---|---|---|---|---|
| 1 | 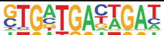 | Tbx20(T-box) | 1e-134 | 3.71% | 56.86% |
| 2 | 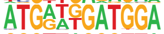 | HOXA1(Homeobox) | 1e-25 | 0.12% | 7.06% |
| 3 | 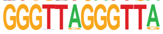 | PSE(SNAPc) | 1e-12 | 0.01% | 2.35% |

**F**

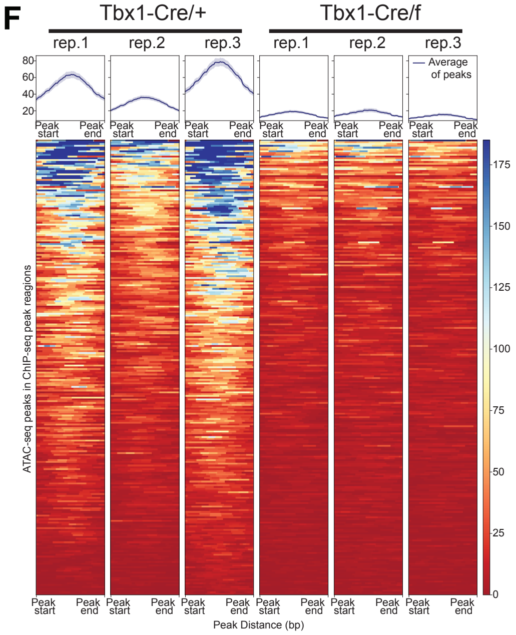
